# BioSensor-X: A Coverage-Aware Evidence-Integration and Benchmarking Framework for Transcription-Factor Prioritization in Microbial Biosensor Engineering

**DOI:** 10.64898/2026.11.22.768397

**Authors:** Mohd Mehboob Uddin, Syed Mohd Zakariya Ali Khan, Sridhar Gunde

## Abstract

Transcription-factor (TF) prioritization in microbial biosensor engineering requires integration of heterogeneous molecular and regulatory evidence, yet benchmark-associated performance may be influenced by candidate coverage, annotation density, and information-flow structure. BioSensor-X was developed as a leakage-controlled, coverage-aware framework for evaluating heterogeneous transcriptomic and regulatory evidence for *Escherichia coli* TF prioritization. Six transcriptomic datasets were processed, of which three contributed candidate instances, yielding 360 candidate contexts representing 60 TFs. The framework integrated transcriptional responsiveness, regulatory specificity, regulon size as an annotation-density proxy, and curated regulatory evidence using discovery-only standardization, prespecified equal weighting, and TF-level maximum aggregation. Performance was evaluated using mutually exclusive TF-level DEVELOPMENT and LOCKED TEST partitions with family-aware stratification, while candidate coverage was assessed separately from conditional ranking performance. Of 12 LOCKED TEST TFs, five were represented in the discovery candidate universe, corresponding to 41.7% coverage. The primary BioSensor-X score achieved a conditional AUROC of 0.5782 (95% bootstrap CI, 0.3308–0.8109), with 1/5 TFs recovered in the Top-10 and 2/5 in the Top-20. The regulon-size/annotation-density component achieved an AUROC of 0.7945 (95% bootstrap CI, 0.6655–0.9055). The integrated framework retained a measurable benchmark-associated ranking signal, while the results also indicated that annotation structure contributed substantially to benchmark discrimination. These findings support the value of separating candidate coverage from conditional ranking performance and explicitly accounting for annotation-density-associated effects when evaluating TF prioritization frameworks.

## 1. Introduction

Microbial biosensors couple a defined molecular or physiological state to a measurable biological output and have become useful in metabolic engineering, strain screening, pathway optimization, and monitoring of cellular states. Transcription factors (TFs) are important components of microbial biosensors because ligand-responsive and stress-responsive regulators can connect intracellular or extracellular signals to transcriptional outputs. TF-based systems therefore provide a means of converting molecular or physiological changes into experimentally measurable reporter signals or engineered pathway responses [19,21,35]. Genetically encoded biosensors encompass additional architectures, including fluorescent protein-based systems and other genetically encoded sensing mechanisms, but TF-mediated designs are particularly relevant when sensing depends on endogenous or engineered transcriptional regulation [6,9,35].

The effectiveness of a TF-based biosensor depends on more than the existence of a regulatory interaction. A useful sensor must couple a defined input to an appropriate regulatory mechanism and generate a measurable response with sufficient specificity and operational performance for the intended application [4,19,35]. These properties are commonly characterized experimentally using reporter assays, dose-response measurements, screening approaches, and pathway-level phenotyping [19,35]. However, the set of experimentally characterized regulators remains much smaller than the regulatory repertoire encoded by microbial genomes. The increasing availability of transcriptomic and regulatory datasets therefore creates an opportunity to evaluate additional TF candidates computationally, while also introducing an evidence-integration problem: datasets capture different biological states, regulatory resources differ in coverage and confidence, and the amount of available information is highly nonuniform across TFs.

Computational analysis of gene-expression and regulatory-network data provides established approaches for estimating TF activity and inferring regulatory relationships. Network-inference methods such as ARACNE, TIGRESS, and tree-based approaches use different statistical assumptions to infer regulatory structure from expression data [2,12,14,20,23]. Benchmarking studies have shown that the performance of regulatory-network inference methods varies with network structure, dataset characteristics, and evaluation design, emphasizing the importance of empirical benchmarking rather than reliance on a single modeling strategy [12,24,29]. Other approaches infer TF-associated activity by combining gene-expression signatures with prior TF-target relationships or network-derived activity estimates [3,10,13,16]. ChEA3 integrates multiple sources of regulatory evidence for TF enrichment, while decoupleR provides an ensemble framework for estimating biological activities from omics data [13,16]. SCENIC and related regulatory-network approaches similarly combine expression information with prior regulatory structure to infer TF-associated activity or regulatory networks [1].

The dependence of TF inference on prior regulatory resources introduces an additional source of variation. Different collections of TF-target relationships can produce different estimates of TF activity, while the coverage and composition of regulons can materially influence the resulting inference [10]. Efforts to expand high-confidence regulon resources have likewise demonstrated that the completeness and reliability of prior regulatory information affect TF-activity estimation and interpretation [22]. These observations become particularly important when computational activity estimates are subsequently converted into TF rankings, because ranking behavior may reflect both biological signal and the amount of prior regulatory annotation available for a given regulator.

This issue becomes more consequential when computational rankings are evaluated against curated biological benchmarks. Such benchmarks are necessarily constructed from available experimental knowledge and may therefore represent the biological space unevenly. Well-characterized TFs can accumulate larger regulons, more experimentally supported regulatory interactions, and greater representation in reference resources than less-characterized regulators. These differences can generate associations between benchmark membership and annotation density that are not necessarily equivalent to the biological property represented by the benchmark endpoint. Large-scale benchmarking of regulatory-network inference has shown that evaluation outcomes depend on benchmark characteristics and underlying network structure [12,24]. In *Escherichia coli*, extensive experimental work has established a substantial transcriptional regulatory network, while resources such as RegulonDB and EcoCyc provide curated representations of this regulatory knowledge [8,17,18,28,30], while the uneven accumulation of such information across regulators also creates the possibility that annotation structure contributes to computational benchmark performance.

This consideration is particularly relevant to computational TF prioritization for microbial biosensor engineering. Regulon size, transcriptional responsiveness, regulatory specificity, and curated regulatory evidence describe different dimensions of the available evidence and should not be assumed to measure the same biological property. Combining these dimensions into a composite score likewise does not guarantee that the resulting ranking will be more informative than any individual component. The behavior of an integrated score depends on the distributions, scaling, dependencies, and benchmark associations of its constituent features. Integration can capture complementary information, but it can also dilute a strong signal carried by a single component. Component-level evaluation is therefore necessary to determine whether evidence integration provides additional benchmark-relevant discrimination.

BioSensor-X was developed to examine this methodological problem in the context of *E. coli* TF prioritization for microbial biosensor applications. The final framework integrates four evidence dimensions: transcriptional responsiveness, regulatory specificity, regulon size treated explicitly as an annotation-density proxy, and curated regulatory evidence. Candidate-level evidence is standardized using parameters derived from the discovery candidate universe, combined using prespecified equal weights, and aggregated to the TF level across candidate contexts. The framework uses a TF-level mutually exclusive DEVELOPMENT/LOCKED TEST evaluation design with family-aware allocation, while candidate coverage is assessed separately from conditional ranking performance.

The primary methodological question was not whether additional evidence dimensions would necessarily improve TF prioritization, but whether heterogeneous evidence integration provides additional benchmark-relevant discrimination after candidate coverage and evidence structure are made explicit. The evaluation therefore compared the integrated score with individual evidence components and prespecified comparator formulations and examined the relationship between regulon size and benchmark-associated signal. Sensitivity and residualized analyses were used to determine whether apparent benchmark-associated signal persisted after accounting for the annotation-density dimension.

Accordingly, this study evaluates the statistical behavior of heterogeneous evidence integration for TF prioritization under explicit coverage and benchmark constraints, with particular emphasis on distinguishing biological evidence from signals arising from the structure of the underlying regulatory annotations. By separating candidate coverage from conditional ranking performance and evaluating individual evidence dimensions alongside the integrated score, BioSensor-X provides a framework for assessing the interpretability and benchmark behavior of TF-prioritization systems before computational scores are treated as proxies for biosensor engineering potential.

## 2. Materials and Methods

### 2.1 Study design, transcriptomic datasets, and information-flow control

BioSensor-X was evaluated using a staged computational design that separated discovery-stage processing from benchmark evaluation. The framework was designed to integrate heterogeneous transcriptomic and regulatory evidence for transcription-factor (TF) prioritization while explicitly separating candidate coverage from conditional ranking performance.

The discovery stage comprised transcriptomic data processing, differential-expression analysis, TF-associated enrichment estimation, candidate construction, evidence-component calculation, discovery-stage standardization, and construction of the primary BioSensor-X score. Benchmark membership was not used to construct the candidate universe, estimate standardization parameters, or define the primary score. Benchmark labels were introduced only after the discovery-stage scoring framework had been specified.

Six *Escherichia coli* transcriptomic datasets were obtained from the NCBI Gene Expression Omnibus (GEO) [5] and processed: GSE4556, GSE4511, GSE12797, GSE20095, GSE34023, and GSE80002. In the final candidate layer, three datasets contributed candidate instances: GSE4556, GSE4511, and GSE12797. The remaining three processed datasets, GSE20095, GSE34023, and GSE80002, contributed no candidate instances to the final candidate matrix. The three contributing datasets generated 360 candidate instances corresponding to 60 TFs represented across six discovery contrasts per TF.

GSE4556 represented anaerobic pH perturbation, GSE4511 represented aerobic pH perturbation, and GSE12797 represented rpoS-associated perturbation under exponential- and stationary-phase conditions. GSE20095, GSE34023, and GSE80002 were processed as part of the broader discovery data layer but yielded no candidate instances under the final candidate-construction rules.

All analysis steps determining the primary candidate-level scores were completed before benchmark labels were used for evaluation. Standardization parameters were estimated from the discovery candidate set and subsequently frozen.

### 2.2 Regulatory resources and candidate-universe construction

The initial regulatory universe comprised E. coli TFs with Tier A or Tier B regulatory annotations and at least three curated regulatory targets. Regulatory target information was derived principally from RegulonDB and EcoCyc [17,28].

A total of 119 TFs satisfied the regulatory criterion of Tier A+B annotation with at least three targets. The final candidate universe was then restricted to TFs for which corresponding transcriptomic evidence was available in the processed discovery data, resulting in 60 TFs. Candidate-universe construction was benchmark-independent; benchmark membership was not used as an inclusion or exclusion criterion.

For TF *t*, the curated regulatory target set was defined as

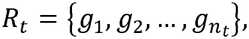

where *n*_t_ ≥ 3 denotes the number of eligible Tier A+B regulatory targets associated with TF *t*. The canonical Tier A+B target count defined here was used consistently for regulon-size scoring, annotation-density diagnostics, matched analyses, and comparator evaluation.

Each candidate instance was defined by a TF and a discovery contrast. Thus, for candidate *i*,

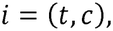

where *t* denotes the TF and *c* denotes the discovery contrast. The final candidate layer contained 360 candidate instances across 60 TFs, with six candidate contexts per TF.

TF-name normalization was applied before benchmark matching to reconcile predefined aliases, including Fnr/FNR and HNS/H-NS.

### 2.3 Differential-expression analysis and TF-activity estimation

Gene-level differential-expression analyses were performed separately for each discovery contrast. limma moderated *t*-statistics were used as the gene-level statistics for downstream TF-associated enrichment analysis [26].

For TF *t* and discovery contrast *c*, let *R*_t_ denote the corresponding set of curated Tier A+B target genes, with

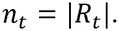

Let *t_g,c_* denote the limma moderated *t*-statistic for gene *g* in contrast *c*. The observed TF-associated enrichment statistic was defined as the mean moderated *t*-statistic across the TF target set:

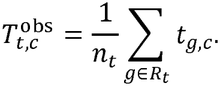

For each TF-contrast pair, a null distribution was generated by randomly sampling *n*_*t*_ genes from the complete set of gene-level moderated *t*-statistics for the corresponding contrast. For permutation *b*= 1,…,*B*, the null statistic was calculated as

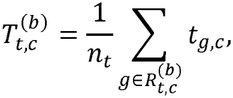

where 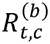 is a randomly sampled set of *n_t_* genes and *B* = 1000.

The mean and standard deviation of the resulting null distribution were calculated as

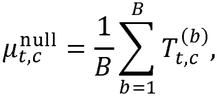

and

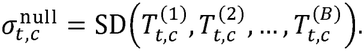

The TF-associated enrichment Z-score was then defined as

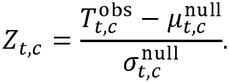

For reproducibility, this quantity is stored as mlm_z in the repository and source data files. The mlm_z field is a legacy variable name and does not denote a mixed-effects linear model or a mixed-effects-model Z-score; it represents the permutation-enrichment Z-score defined above. The analysis therefore used permutation-based enrichment of limma moderated t-statistics. The implementation details are documented in the reproducibility package.

The corresponding two-sided empirical P value was calculated as

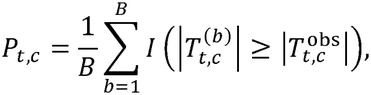

where *I*(·) is the indicator function. Raw empirical P values were retained without multiple-testing adjustment in the candidate-level data layer.

For downstream scoring, the transcriptional-responsiveness component for candidate *i*, corresponding to TF *t* and contrast *c*, was defined as the absolute TF-associated enrichment score:

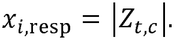

This formulation permits both positive and negative regulatory responses to contribute to the responsiveness magnitude without treating the direction of regulation itself as evidence of biosensor suitability.

### 2.4 Evidence-component construction

The final BioSensor-X score comprised four evidence dimensions:

1. Transcriptional Responsiveness;
2. Regulatory Specificity;
3. Regulon size as an annotation-density proxy
4. Curated Regulatory Evidence.

These components were intended to represent different aspects of the available transcriptomic and regulatory evidence rather than equivalent biological quantities. In particular, regulon size was treated throughout the analysis as an annotation-density proxy rather than as a direct measurement of mechanistic depth or experimentally demonstrated biosensor suitability.

#### 2.4.1 Transcriptional responsiveness

For candidate *i*,

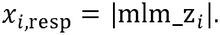

The absolute magnitude was used because the direction of TF-associated transcriptional change can depend on the intended biosensor architecture and does not, by itself, establish biosensor suitability.

#### 2.4.2 Regulatory specificity

For each candidate *i*, the target-context response was compared with the distribution of responses for the same TF in other available signal contexts. Let

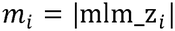

denote the absolute target-context TF-associated activity, and let

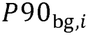

denote the 90th percentile of absolute mlm_z values for the same TF across its other available contexts.

The raw specificity component was

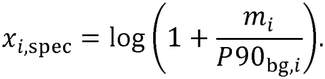

Use of the 90th percentile rather than an extreme maximum reduces sensitivity to a single unusually large background response.

#### 2.4.3 Regulon size and annotation density

For TF *t*, the regulon-size component was calculated as

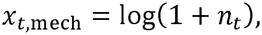

where *n_t_* is the canonical number of eligible Tier A+B curated regulatory targets defined in Section 2.2.

Because this quantity is a monotonic transformation of curated target count, it was interpreted as a proxy for annotation density throughout the analysis rather than as a direct measure of mechanistic depth or biosensor performance.

#### 2.4.4 Curated regulatory evidence

Regulatory evidence was mapped from the curated regulatory-resource classification to a predefined ordinal scale:

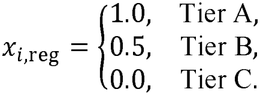

The regulatory evidence component therefore encoded the relative strength of available curated regulatory support rather than direct biosensor-performance measurements.

### 2.5 Discovery-only standardization and primary score construction

Each raw evidence component was standardized using the complete 360-candidate discovery set. For component *k*, with raw value *x*_i,k_, the discovery mean was

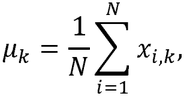

where *N* = 360, and the corresponding standard deviation was

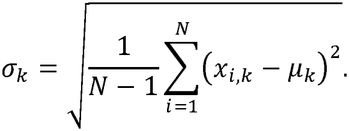

The standardized component was

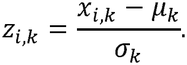

The resulting means and standard deviations were frozen before benchmark evaluation and applied unchanged thereafter. No benchmark labels were used during standardization. The mechanistic component was calculated from the canonical Tier A+B target count defined in Section 2.2.

The four standardized components were combined using prespecified equal weights:

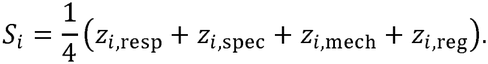

Equivalently,

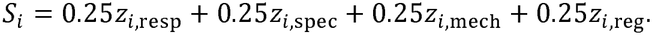

The equal weights were fixed independently of LOCKED TEST performance. Benchmark membership was excluded from the primary score.

### 2.6 Candidate-to-TF aggregation and benchmark construction

Because each TF was represented by multiple candidate contexts, candidate-level scores were converted to TF-level scores using maximum aggregation.

For TF *t*, with candidate set *I_t_*,

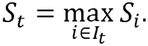

The candidate attaining the maximum score was retained as the best-scoring context for that TF. For tied primary scores, the implemented procedure used the candidate with the largest absolute mlm_z value as the tie-breaker. The same aggregate_tf() maximum-based procedure was used for the applicable comparator calculations.

The mean candidate-level score was retained descriptively but was not used for the primary TF ranking. Maximum aggregation was the prespecified TF-level aggregation strategy used for the primary analysis.

The curated benchmark comprised 112 E. coli TF-associated systems. At the final TF-level evaluation stage, the benchmark was divided into mutually exclusive DEVELOPMENT and LOCKED TEST sets. The corrected partition contained 29 DEVELOPMENT TFs and 12 LOCKED TEST TFs, with no TF occurring in both partitions. The allocation was family-aware/stratified at the TF level rather than a strict family-disjoint holdout.

#### The LOCKED TEST TF set was

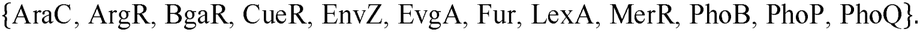

Candidate representation was assessed independently of benchmark performance. A benchmark TF was considered covered when at least one candidate instance for that TF occurred in the 60-TF discovery candidate universe.

#### The five represented LOCKED TEST TFs were

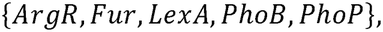

while AraC, BgaR, CueR, EnvZ, EvgA, MerR, and PhoQ had no candidate representation. Thus,

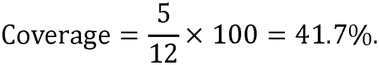

### 2.7 Primary performance evaluation and statistical analysis

The primary endpoint was conditional AUROC on the corrected LOCKED TEST benchmark. Conditional evaluation was restricted to TFs represented in the discovery candidate universe. This yielded five positive TFs and 55 negative TFs within the fixed 60-TF candidate population. The positive class comprised the five covered LOCKED TEST TFs; the negative class comprised all remaining candidate TFs that were not members of the corrected LOCKED TEST set.

AUROC was used as the primary discrimination measure [11]. For TF-level ranking, Top-5, Top-10, and Top-20 recovery were calculated. A benchmark TF was considered recovered at Top-*k* when its TF-level rank satisfied

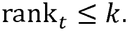

Mean reciprocal rank was calculated as

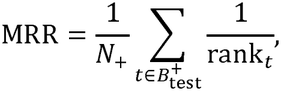

where *N*_+_ is the number of represented positive TFs.

Median rank was calculated across the represented positive TFs.

### Bootstrap confidence intervals

Uncertainty in AUROC was quantified using 10,000 TF-level bootstrap replicates. In each replicate, the five positive TFs and 55 negative TFs were independently resampled with replacement while preserving the original class sizes:

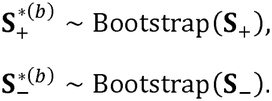

For bootstrap replicate *b*,

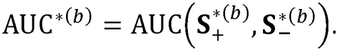

The 95% percentile confidence interval was defined as

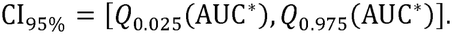

The bootstrap estimates were independently cross-checked against analytical Hanley-McNeil AUROC intervals [7,11]. The final primary estimate was AUROC 0.5782 with a bootstrap 95% confidence interval of 0.3308–0.8109. The corresponding Top-5, Top-10, and Top-20 recovery rates were 1/5 (20.0%), 1/5 (20.0%), and 2/5 (40.0%), respectively. The mean reciprocal rank was 0.0693, with a median rank of 22.

Because five positive TFs were represented in the LOCKED TEST candidate universe, uncertainty was assessed at the TF level using bootstrap confidence intervals together with ranking and coverage metrics. The results were interpreted using these complementary measures rather than AUROC alone.

### 2.8 Comparator analyses, annotation-density diagnostics, reproducibility, and data/code availability

The comparator set comprised the predefined benchmark and evidence-based comparators retained for the final evaluation, including expression-based, regulatory, TF-activity, benchmark-membership, and annotation-density controls.

#### Expression Only (B1)

B1 was defined as an expression-based comparator using the absolute standardized transcriptional-responsiveness component alone:

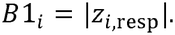

#### Regulatory Only (B2)

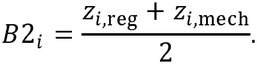

B2 was defined as a regulatory-evidence comparator combining the standardized curated regulatory-evidence and regulon-size components with equal weighting.

#### Custom TF-Activity Proxy (B4)

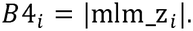

B4 was defined as a custom permutation-enrichment TF-activity proxy. It was evaluated as a diagnostic evidence-based comparator distinct from the primary BioSensor-X score.

#### Benchmark-Information Control (B7)

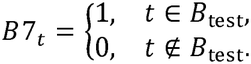

B7 was defined as a benchmark-union membership control, assigning a value of 1 to TFs occurring in either the DEVELOPMENT or LOCKED TEST partitions and 0 otherwise. This comparator was included as a benchmark-information control to characterize the extent to which broad prior benchmark membership contributes to discrimination within the evaluated candidate universe.

#### Mechanistic/annotation-density component (B8)

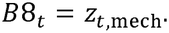

B8 was defined as the standardized regulon-size component based on canonical Tier A+B target count and was used as an annotation-density comparator.

#### Annotation-density diagnostics

To examine the relationship between benchmark-associated discrimination and annotation structure, TFs were stratified by canonical regulon size and component performance was compared across annotation-density strata. Residualized analyses were additionally used to assess the extent to which the apparent benchmark-associated signal of other evidence dimensions remained after accounting for the annotation-density proxy.

Because the annotation-density component is defined directly from curated target count, its association with regulon size is expected by construction. These analyses were therefore interpreted as diagnostics of feature structure rather than as evidence of a causal relationship between annotation density and biosensor suitability.

All sensitivity analyses were conducted using the DEVELOPMENT partition and did not alter the LOCKED TEST endpoint or the frozen primary score.

#### Information-leakage controls

The following safeguards were maintained throughout the analysis:

**candidate construction → discovery-only feature calculation → frozen standardization → frozen score construction → benchmark evaluation**.

Benchmark labels were not used to construct the candidate universe or estimate discovery-stage standardization parameters. Equal component weights were prespecified, and LOCKED TEST outcomes were not used to tune the primary score or its aggregation strategy.

The benchmark-membership feature was excluded from the primary BioSensor-X score and was used only for the B7 positive-control analysis.

#### Computational implementation and reproducibility

Statistical analyses were implemented in R [25], with complementary workflow and data-processing components implemented in Python. Candidate-level and TF-level outputs, benchmark files, frozen standardization parameters, derived tables, and figure-generation materials were retained as fixed reproducibility artifacts.

The final reproducibility package contains the source code, processed candidate data, benchmark definitions, standardization parameters, analysis outputs, and supplementary-generation resources required to reproduce the reported workflow. The public BioSensor-X repository serves as the repository of record for the computational materials described in this study.

## 3. Results

### 3.1 Candidate universe, benchmark composition, and candidate coverage

The discovery-stage regulatory filtering identified 119 E. coli transcription factors with Tier A or Tier B regulatory annotations and at least three eligible regulatory targets. Restriction to TFs with corresponding transcriptomic evidence in the processed discovery data yielded a final candidate universe of 60 TFs. Three of the six processed transcriptomic datasets contributed candidate instances to this universe: GSE4556, GSE4511, and GSE12797. These datasets contributed six discovery contrasts per TF, resulting in 360 candidate instances. GSE20095, GSE34023, and GSE80002 were processed as part of the discovery data layer but generated no candidate instances under the final candidate-construction criteria.

The curated benchmark contained 112 E. coli TF-associated systems, comprising 25 Tier 1 entries corresponding to demonstrated biosensor systems and 87 Tier 2 entries corresponding to experimentally supported TF-regulated systems relevant to biosensor development. The corrected benchmark was partitioned at the TF level into mutually exclusive DEVELOPMENT and LOCKED TEST sets containing 29 and 12 TFs, respectively, with no TF overlap. The allocation was family-aware and stratified rather than a strict family-disjoint holdout.

Candidate coverage was assessed against the LOCKED TEST benchmark to determine the extent to which benchmark TFs were represented in the discovery candidate universe. Five of the 12 LOCKED TEST TFs were represented: ArgR, Fur, LexA, PhoB, and PhoP. The remaining seven TFs were AraC, BgaR, CueR, EnvZ, EvgA, MerR, and PhoQ. Overall candidate coverage was therefore 5/12, corresponding to 41.7%.

**Table 1.** Study design, candidate universe, benchmark composition, and LOCKED TEST candidate coverage.

| Aspect | Detail |
|---|---|
| **Candidate Universe** |  |
| Discovery datasets (GEO) | GSE4511, GSE4556, GSE12797, GSE20095, GSE34023, GSE80002 |
| Number of datasets | 6 |
| Signal contexts | 6 (high/low pH, aerobic/anaerobic; rpoS-associated exponential/stationary) |
| Total candidate instances | 360 |
| Unique TFs represented | 60 |
| Candidates per TF | 6 (one per signal context) |
| **Benchmark** |  |
| Total curated TF-associated systems | 112 |
| Tier 1 (demonstrated biosensor systems) | 25 |
| Tier 2 (experimentally supported TF-regulated systems relevant to biosensor development) | 87 |
| Partition method | 70/30 TF-level allocation with family-aware/stratified assignment |
| **DEVELOPMENT** |  |
| DEVELOPMENT benchmark TFs | 29 |
| Benchmark TFs with candidate representation | 13/29 (44.8%) |
| Benchmark TFs without candidate representation | 16/29 (55.2%) |
| **LOCKED TEST** |  |
| LOCKED TEST benchmark TFs | 12 |
| Benchmark TFs with candidate representation | 5/12 (41.7%) |
| Represented LOCKED TEST TFs | ArgR, Fur, LexA, PhoB, PhoP |
| Benchmark TFs without candidate representation | 7/12 (58.3%) |
| Unrepresented LOCKED TEST TFs | AraC, BgaR, CueR, EnvZ, EvgA, MerR, PhoQ |
| Candidate-coverage interpretation | Coverage was assessed separately from conditional ranking performance; conditional ranking was evaluated within the represented candidate population. |

Candidate coverage and conditional ranking performance were therefore treated as distinct evaluation properties. The conditional LOCKED TEST analysis evaluated ranking behavior within the represented candidate population, while candidate coverage quantified the extent of benchmark representation achieved by the discovery-stage candidate construction.

### 3.2 Primary performance on the LOCKED TEST benchmark

The primary endpoint was conditional AUROC on the corrected LOCKED TEST benchmark. Within the fixed 60-TF candidate universe, five represented LOCKED TEST TFs constituted the positive class and the remaining 55 candidate TFs constituted the negative class.

The equal-weight BioSensor-X Primary score achieved an AUROC of 0.5782 with a 95% TF-level bootstrap confidence interval of 0.3308–0.8109. The corresponding recovery rates were 1/5 (20%) at Top-5, 1/5 (20%) at Top-10, and 2/5 (40%) at Top-20. The mean reciprocal rank was 0.0693, and the median rank of the represented benchmark-positive TFs was 22.0.

The bootstrap interval provides an explicit measure of uncertainty around the TF-level estimate, while the Top-k recovery, mean reciprocal rank, median rank, and candidate-coverage measures provide complementary information on ranking behavior.

The primary score produced a measurable benchmark-associated ranking signal within the represented LOCKED TEST population. Candidate coverage and conditional ranking were evaluated as distinct properties of the framework, with ranking performance assessed within the represented candidate population.

**Table 2.** Conditional LOCKED TEST performance of the BioSensor-X Primary score and prespecified comparator methods.

| **Method** | **AUROC (95% bootstrap CI)** | **Top-5** | **Top-10** | **Top-20** | **Role** |
|---|---|---|---|---|---|
| BioSensor-X Primary | 0.5782 (0.3308–0.8109) | 1/5 (20%) | 1/5 (20%) | 2/5 (40%) | Primary |
| B1 Expression Only | 0.2945 (0.1127–0.4909) | 0/5 (0%) | 0/5 (0%) | 0/5 (0%) | Comparator |
| B2 Regulatory Only | 0.6545 (0.3818–0.9091) | 1/5 (20%) | 2/5 (40%) | 3/5 (60%) | Comparator |
| B4 Custom TF-Activity Proxy | 0.3527 (0.1964–0.5236) | 0/5 (0%) | 0/5 (0%) | 0/5 (0%) | Comparator |
| B7 Benchmark-Information Control | 0.8818 | 0/5 (0%) | 5/5 (100%) | 5/5 (100%) | Benchmark-information control |
| B8 Annotation-Density Diagnostic | 0.7945 (0.6655–0.9055) | 1/5 (20%) | 1/5 (20%) | 5/5 (100%) | Annotation-density diagnostic |

### 3.3 Component-level and comparator performance

The individual evidence dimensions showed distinct benchmark-associated behavior on the corrected LOCKED TEST population. The regulon-size component, designated B8 in the evaluation framework, achieved an AUROC of 0.7945 with a 95% bootstrap confidence interval of 0.6655–0.9055. B8 recovered 1/5 (20%) of represented positives in the Top-5, 1/5 (20%) in the Top-10, and 5/5 (100%) in the Top-20. The regulatory-only comparator B2 achieved an AUROC of 0.6545 (95% bootstrap CI, 0.3818–0.9091), with 1/5, 2/5, and 3/5 represented positives recovered at Top-5, Top-10, and Top-20, respectively. The expression-only comparator B1 and custom TF-activity proxy B4 provided additional single-axis reference points for comparison with the integrated score. An annotation-matched paired comparison across five TF pairs provided an additional ranking-based assessment. BioSensor-X ranked higher in one pair and lower in four pairs (paired win rate = 0.2000); the Wilcoxon signed-rank test was not significant (*p* = 0.90625) [33].

The benchmark-information control B7 achieved an AUROC of 0.8818. Because B7 was defined from prior membership in the DEVELOPMENT or LOCKED TEST benchmark partitions, it provides a benchmark-information reference rather than an independent evidence-based predictor.

Taken together, the comparator results indicate that the four-component BioSensor-X score did not exceed the strongest individual component on the evaluated benchmark. The BioSensor-X Primary AUROC of 0.5782 was lower than the B8 regulon-size/annotation-density diagnostic at 0.7945. This comparison reflects the behavior of the evaluated score within the present benchmark structure and does not imply that larger regulons are intrinsically superior biosensor candidates.

### 3.4 Regulon size and annotation-density-associated signal

The benchmark-associated discrimination of B8 was examined in relation to the underlying regulon-size variable from which the component was derived. B8 is calculated from curated regulatory target count and is therefore a monotonic transformation of regulon size. The resulting association with regulon size is expected from the definition of the component and was evaluated as an annotation-density characteristic rather than as an independent measure of biosensor performance.

The relationship between regulon size and benchmark representation was therefore examined directly. Figure 4 compares the regulon-size distribution of the represented LOCKED TEST TFs with that of the remaining candidate TFs in the 60-TF candidate universe. The comparison provides context for interpreting the benchmark-associated performance of B8 and indicates that part of the observed signal is associated with differences in the density of curated regulatory annotation.

**Figure 1.**
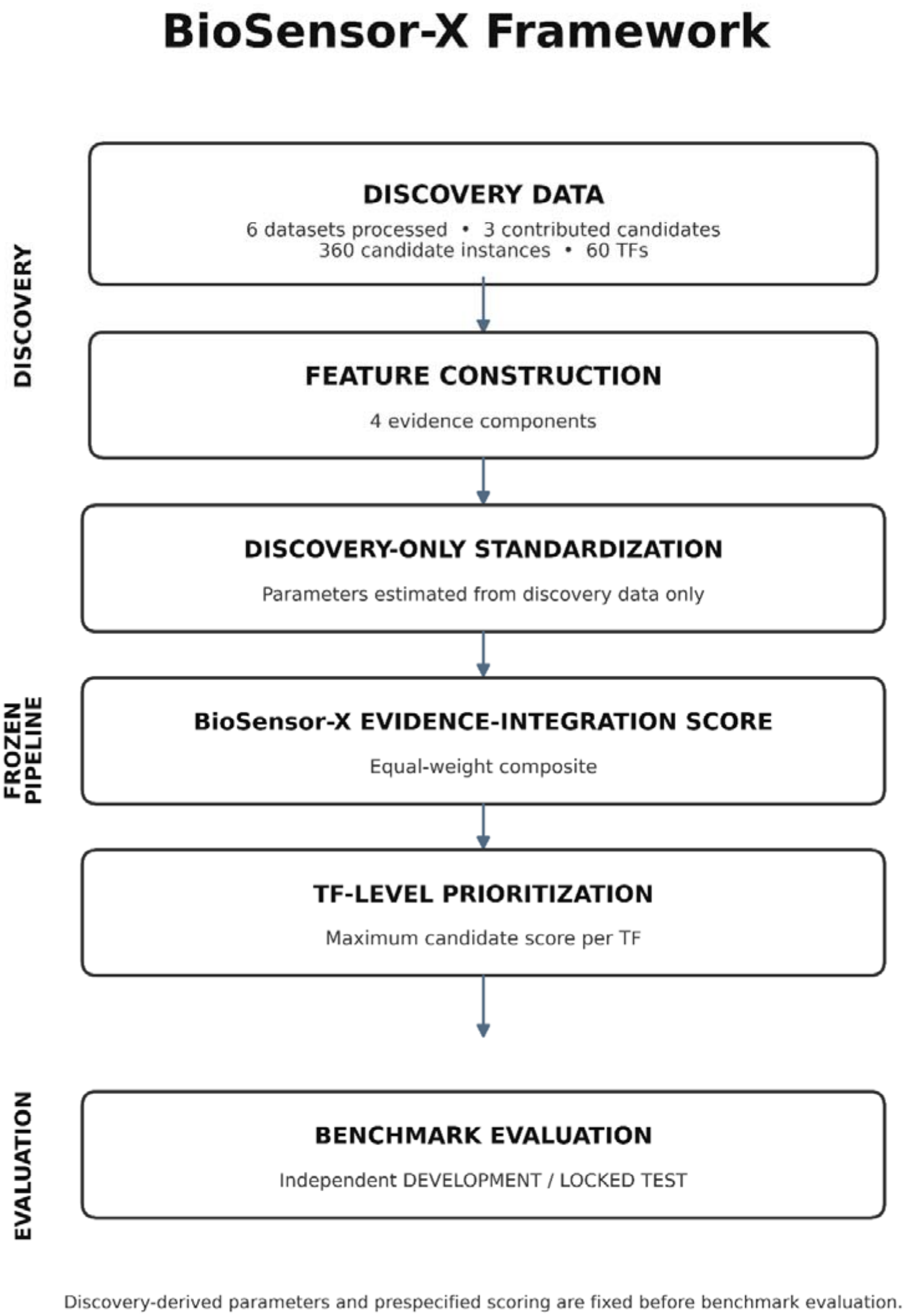
BioSensor-X computational workflow.

**Figure 2.**
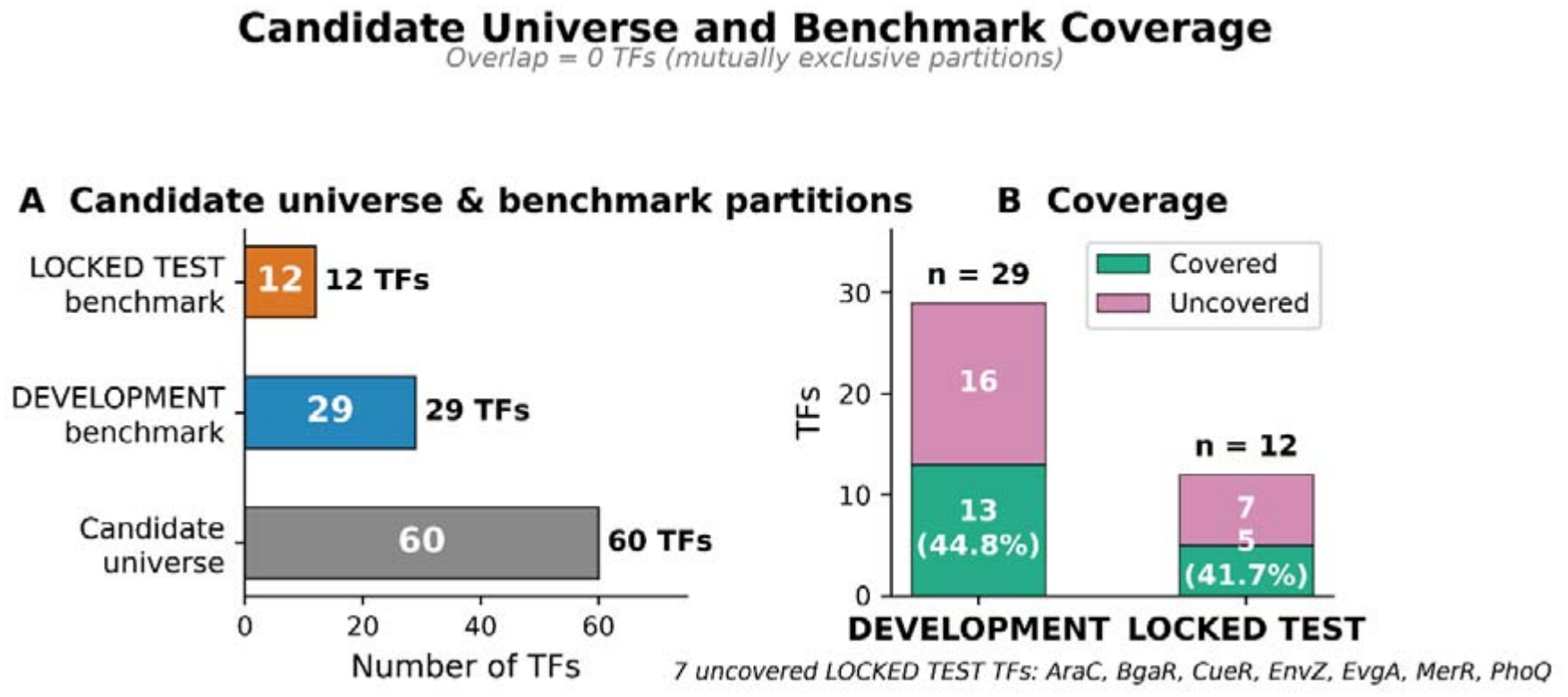
Candidate representation and coverage across the benchmark.

**Figure 3.**
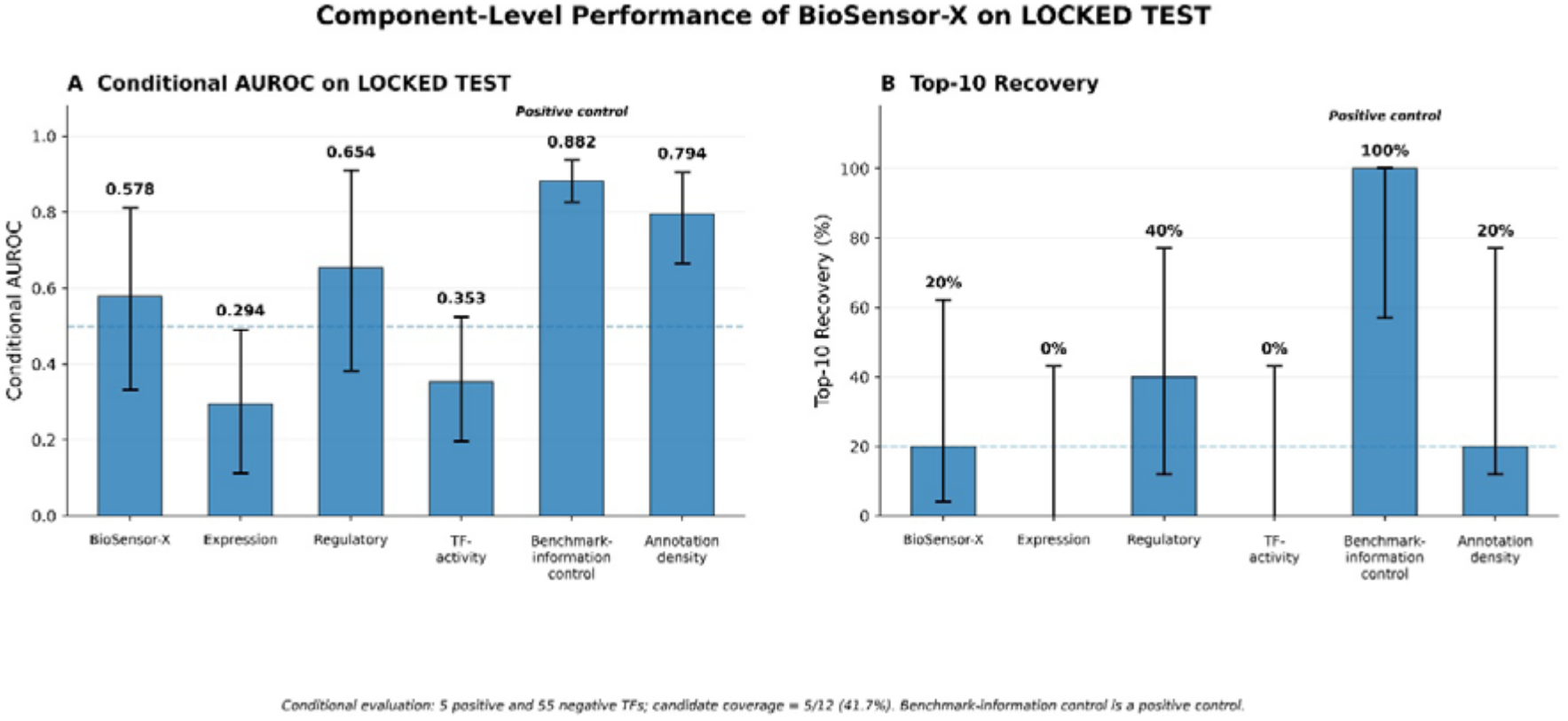
Component-level and comparator performance on the corrected LOCKED TEST evaluation.

**Figure 4.**
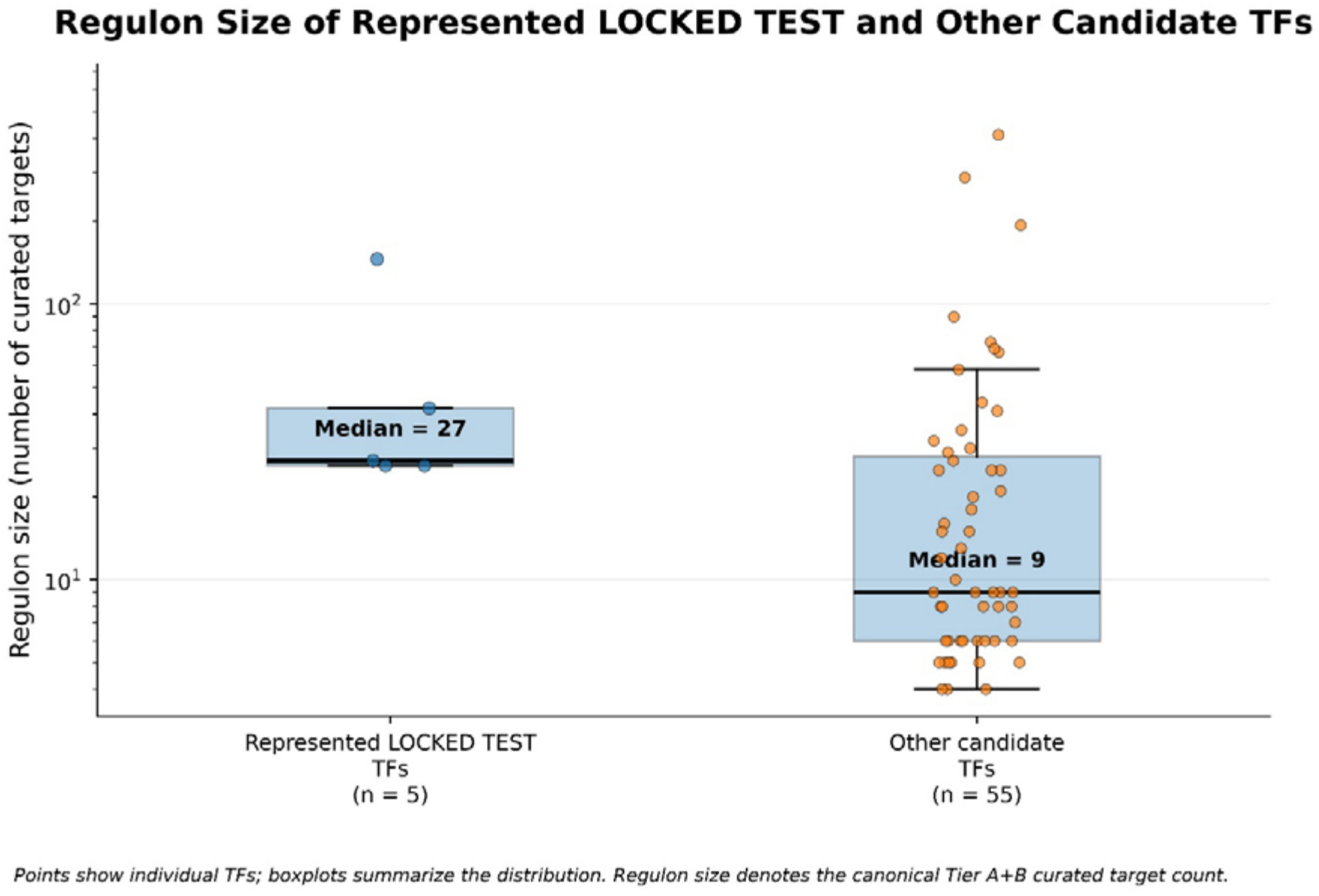
Regulon-size distributions of represented LOCKED TEST TFs and other candidate TFs. Regulon size is defined as the canonical number of eligible Tier A+B curated regulatory targets specified in Section 2.2. The figure compares the five represented LOCKED TEST TFs with the remaining 55 candidate TFs.

To examine the annotation structure further, the 60-TF candidate universe was ordered by canonical Tier A+B target count, with TF name used to resolve ties, and divided into three equal rank-based strata of 20 TFs each. The resulting observed target-count ranges were 4–8 for the low stratum, 9–25 for the medium stratum, and 26–413 for the high stratum. Figure 5 summarizes the distribution of DEVELOPMENT benchmark TFs across these annotation-density strata.

**Figure 5.**
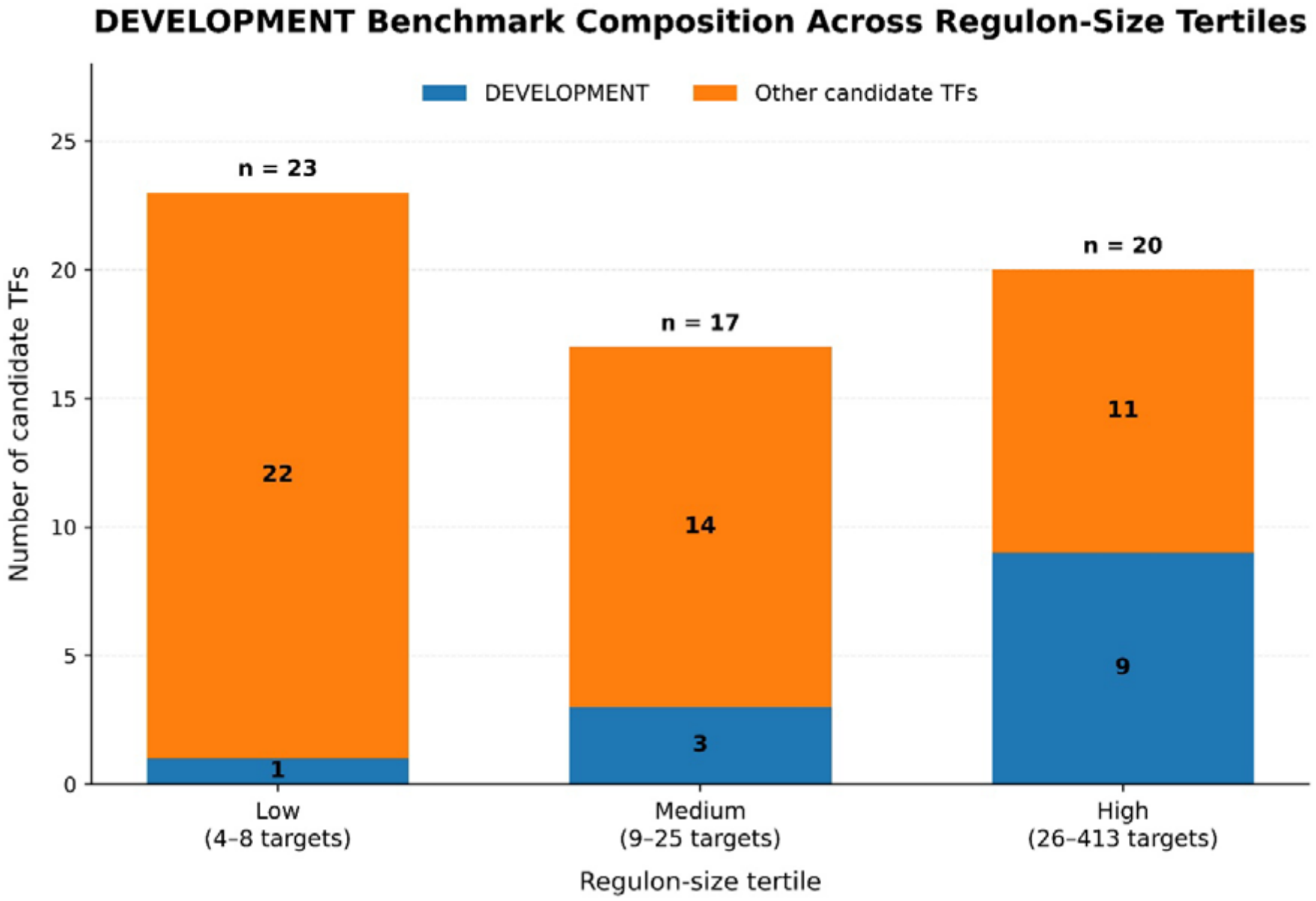
DEVELOPMENT benchmark composition across regulon-size tertiles. The 60-TF candidate universe was ordered by canonical Tier A+B target count, with alphabetical TF-name ordering used to resolve ties, and divided into three equal rank-based strata of 20 TFs each. The observed target-count ranges were 4–8, 9–25, and 26–413 for the low, medium, and high strata, respectively. The figure summarizes the composition of the DEVELOPMENT partition across these strata.

**Figure 6.**
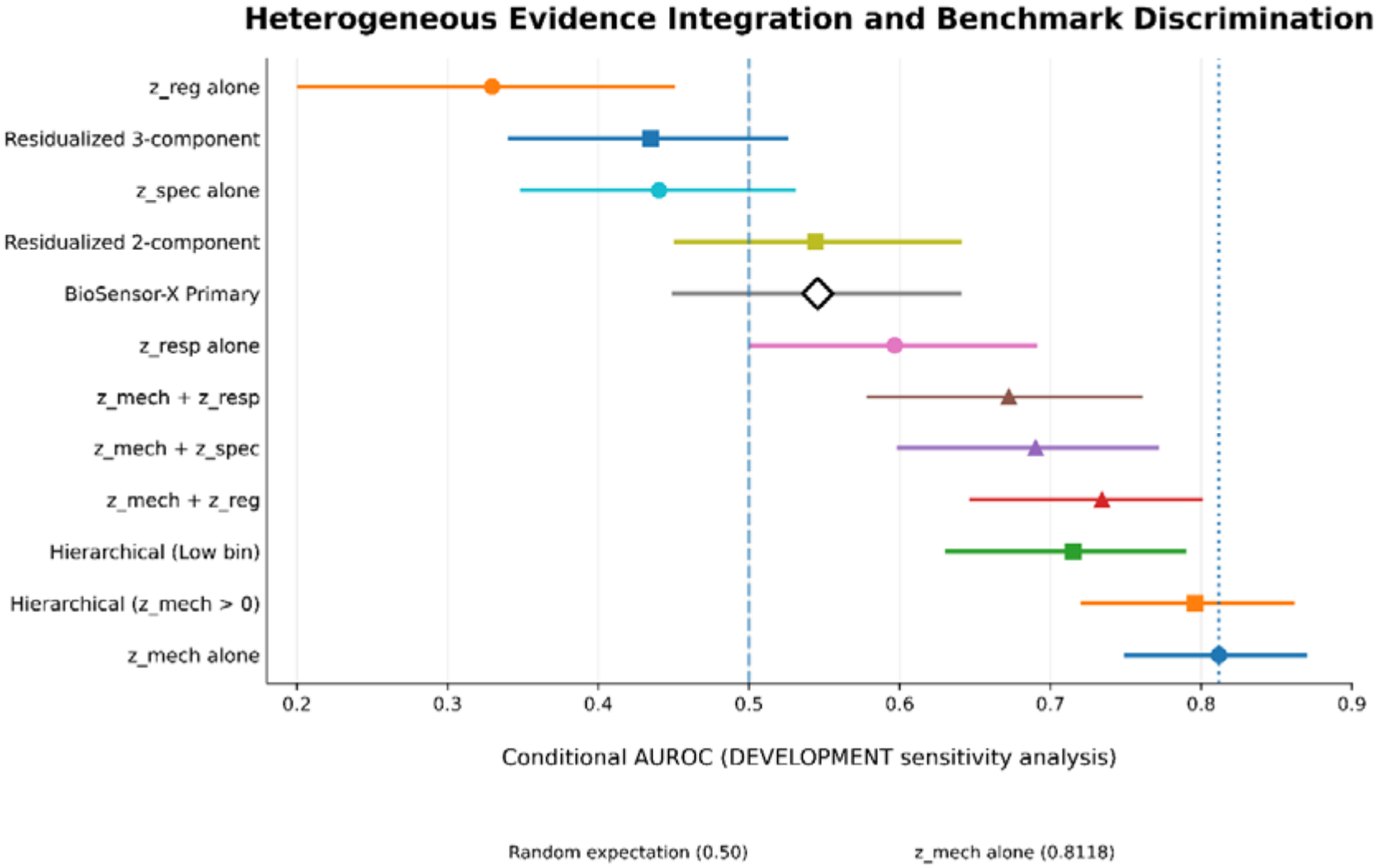
DEVELOPMENT-partition performance of the BioSensor-X primary composite and diagnostic integration schemes.

Together, Figures 4 and 5 place the B8 result in the context of the underlying annotation structure. The distribution of benchmark representation across annotation-density strata supports interpretation of B8 as an annotation-density-associated diagnostic rather than as a direct measure of mechanistic superiority or biosensor suitability.

### 3.5 Residualized and sensitivity analyses

Residualization analyses were used to examine the extent to which benchmark-associated signal remained after accounting for the regulon-size/annotation-density dimension. The canonical residualized analysis yielded an AUROC of 0.4910 for the primary score, indicating that adjustment for annotation density did not materially strengthen the integrated ranking signal. Complementary residualized component and component-subset analyses were conducted within the DEVELOPMENT partition to characterize the behavior of the individual evidence dimensions and alternative combinations.

The sensitivity analyses were used to examine whether the observed ranking behavior was dependent on a particular evidence combination or integration structure. Across these analyses, the annotation-density component remained an important reference dimension, while alternative combinations provided distinct patterns of benchmark-associated discrimination. These analyses were diagnostic and did not alter the prespecified primary score or the LOCKED TEST endpoint.

Detailed residualized, component-subset, hierarchical, and regulon-size-stratified results are provided in Supplementary Table S3.

### 3.6 Composite integration and DEVELOPMENT diagnostic analyses

The DEVELOPMENT analyses were used to characterize the behavior of the BioSensor-X evidence dimensions under alternative component combinations and diagnostic formulations. The analyses considered the integrated score alongside individual evidence components, residualized formulations, and hierarchical combinations, providing a broader assessment of how the evidence dimensions contributed to benchmark-associated ranking behavior.

The annotation-density component remained an important reference dimension across the DEVELOPMENT analyses, while the integrated formulations produced distinct ranking patterns. The canonical residualized primary analysis yielded an AUROC of 0.4910, indicating that adjustment for the annotation-density dimension did not increase the benchmark-associated discrimination of the integrated score.

These DEVELOPMENT analyses were used for methodological characterization and sensitivity assessment and were kept separate from the LOCKED TEST endpoint. The detailed component-subset, residualized, and hierarchical results are provided in Supplementary Table S3.

### 3.7 Information-flow controls, candidate-coverage constraints, and robustness

The final evaluation incorporated explicit information-flow controls throughout the analysis. The candidate universe was constructed independently of benchmark membership, evidence components were calculated without benchmark labels, and standardization parameters were estimated from the 360 discovery candidates and frozen before evaluation. Equal component weights and the primary aggregation strategy were specified independently of LOCKED TEST performance. Benchmark membership was excluded from the primary BioSensor-X score and used only for the B7 benchmark-information control.

The corrected TF-level benchmark partition contained zero TF overlap between DEVELOPMENT and LOCKED TEST. Five of the 12 LOCKED TEST TFs were represented in the discovery candidate universe, and conditional ranking was evaluated within the resulting 60-TF candidate population. Candidate coverage and conditional ranking were therefore treated as distinct evaluation properties, allowing benchmark representation to be characterized separately from ranking behavior.

Uncertainty in the conditional evaluation was quantified using TF-level bootstrap resampling, with AUROC interpreted together with Top-k recovery, mean reciprocal rank, median rank, and candidate coverage. The final primary AUROC was 0.5782 with a 95% bootstrap confidence interval of 0.3308–0.8109, providing a direct measure of uncertainty around the TF-level estimate.

The candidate-coverage analysis defines the scope of the conditional benchmark evaluation. The reported ranking metrics therefore describe the represented LOCKED TEST population within the predefined discovery candidate universe, while the full benchmark composition is captured separately through the coverage analysis in Section 3.1.

Overall, the corrected evaluation showed that BioSensor-X captured a measurable benchmark-associated ranking signal within the represented LOCKED TEST population, while the regulon-size/annotation-density component provided a stronger reference signal on the same endpoint. The DEVELOPMENT diagnostics further characterized the relationship among the evidence dimensions and the contribution of annotation structure to benchmark-associated ranking behavior. Together, these results support BioSensor-X as a structured framework for integrating and evaluating heterogeneous evidence in TF prioritization for microbial biosensor development.

## 4. Discussion

### 4.1 Principal findings and interpretation

This study evaluated whether integrating heterogeneous transcriptomic and regulatory evidence improves transcription-factor (TF) prioritization when candidate coverage and benchmark structure are explicitly considered. BioSensor-X combines transcriptional responsiveness, regulatory specificity, regulon size as an annotation-density proxy, and curated regulatory evidence using prespecified equal weights. The corrected LOCKED TEST evaluation contained five represented benchmark-positive TFs and 55 negative TFs within the fixed 60-TF candidate universe. The primary BioSensor-X score achieved a conditional AUROC of 0.5782 (95% TF-level bootstrap CI, 0.3308–0.8109), with 1/5 represented positives recovered in the Top-5, 1/5 in the Top-10, and 2/5 in the Top-20. The mean reciprocal rank was 0.0693 and the median rank of the represented benchmark-positive TFs was 22.0. These results indicate a measurable benchmark-associated ranking signal, but also substantial uncertainty in the conditional estimate.

The principal finding was that the integrated four-component score did not exceed the strongest individual evidence dimension. The regulon-size component, represented by B8 as the annotation-density comparator, achieved an AUROC of 0.7945 (95% bootstrap CI, 0.6655–0.9055), which was higher than the 0.5782 obtained by the composite score. This comparison indicates that combining additional evidence dimensions did not produce stronger benchmark-associated discrimination under the present implementation and benchmark structure.

This observation is consistent with the broader computational-biology literature, where the performance of regulatory-network inference and TF-activity estimation depends on the characteristics of the input data, prior regulatory resources, and benchmark construction [10,12,24]. Methods such as ARACNE, TIGRESS, and tree-based regulatory inference make different assumptions about network structure and statistical dependence [12,14,20], while integrative TF-activity frameworks show that the choice and coverage of prior regulatory information can materially influence inferred activity estimates [1,10,13,16,22].

The stronger performance of the regulon-size component should not, however, be interpreted as evidence that TFs with larger regulons are intrinsically better biosensor candidates. In BioSensor-X, regulon size is derived from curated regulatory annotation and is therefore partly a measure of how extensively a TF has been characterized. The observed benchmark association is consequently more appropriately interpreted as an annotation-density-associated signal than as evidence of superior biosensor functionality.

### 4.2 Evidence integration and annotation-density-associated signal

The four BioSensor-X components were intended to capture distinct dimensions of computational evidence rather than interchangeable measurements of biosensor performance. Transcriptomic responsiveness captures condition-associated regulatory activity, regulatory specificity describes the relationship between a target context and other contexts for the same TF, regulon size reflects the extent of curated regulatory annotation, and curated regulatory evidence captures the strength of available regulatory support. This design is conceptually related to existing approaches that combine expression data with prior TF-target information to infer regulatory activity [1,10,13,16,22].

The present evaluation shows that evidence aggregation does not necessarily produce complementary predictive information. The equal-weight composite performed below the regulon-size reference, indicating that combining heterogeneous evidence did not, in this evaluation, produce stronger benchmark-associated discrimination than the strongest individual component. Standardization places heterogeneous features on a comparable scale, but it does not remove correlation, redundancy, or benchmark-specific associations between individual evidence dimensions and the evaluation endpoint.

The behavior of the regulon-size component is particularly important because its mathematical construction directly links the feature to the number of curated regulatory targets. Resources such as RegulonDB and EcoCyc contain substantially richer experimental annotation for some TFs than for others, producing an uneven distribution of regulatory knowledge across the network [17,22,28,30,31]. A literature-derived benchmark can therefore inherit part of this underlying annotation structure. The AUROC of 0.7945 observed for the annotation-density comparator is consistent with such benchmark-associated structure, but it does not establish that annotation density itself determines biosensor suitability.

The relationship between annotation structure and benchmark association was further examined through regulon-size distributions, rank-based annotation-density strata, and residualized analyses. These analyses indicate that benchmark-associated signal is not fully separable from the structure of the curated regulatory resource. The canonical residualized primary analysis yielded an AUROC of 0.4910, indicating that adjustment for the annotation-density dimension did not increase the benchmark-associated discrimination of the integrated score. Together, these findings support interpreting annotation density as an important source of benchmark structure rather than as a biological measure of biosensor performance.

### 4.3 Implications for TF prioritization and benchmark design

Computational TF prioritization is potentially useful for reducing the experimental search space, particularly when large regulatory candidate spaces must be screened. However, a computational ranking system can only prioritize candidates that are represented in the underlying candidate universe. In the present study, 5 of the 12 LOCKED TEST TFs (41.7%) were represented in the discovery candidate universe: ArgR, Fur, LexA, PhoB, and PhoP. The remaining seven TFs, AraC, BgaR, CueR, EnvZ, EvgA, MerR, and PhoQ, were not represented and therefore did not contribute to the conditional ranking analysis.

This distinction between coverage and conditional discrimination is important when interpreting computational prioritization benchmarks. A ranking method cannot recover a candidate that was excluded before ranking begins. Consequently, reporting only AUROC or Top-k recovery can obscure an important upstream limitation of candidate generation. Reporting candidate coverage alongside conditional ranking performance makes these two evaluation properties explicit and prevents the performance of the ranking stage from being interpreted independently of the search space on which it operates.

The coverage pattern also provides biological context for the candidate-generation stage. The uncovered LOCKED TEST TFs include sensor-kinase-associated regulators and regulatory factors with comparatively sparse representation in the available discovery evidence. This suggests that candidate construction based on transcriptomic and curated regulatory resources may preferentially represent better-characterized parts of the regulatory network. For practical biosensor discovery, candidate-generation procedures should therefore be evaluated not only for ranking quality but also for whether they provide adequate representation of the broader biological search space.

The benchmark itself also requires careful interpretation. The final benchmark combines demonstrated biosensor systems with experimentally supported TF-regulated systems relevant to biosensor development. It therefore captures a mixture of established biosensor evidence and broader regulatory relevance rather than a uniform experimental measure of biosensor performance. Established TF-based biosensor studies show that useful biosensor properties include responsiveness, selectivity, dynamic range, and appropriate circuit behavior, characteristics that cannot be inferred solely from regulatory annotation or transcriptomic association [19,27,34,35].

These considerations indicate that benchmark construction should explicitly account for candidate coverage, annotation density, biological context, and endpoint definition. Future benchmarks would benefit from distinguishing experimentally demonstrated biosensor phenotypes from broader regulatory evidence and from clearly defining whether the intended endpoint represents biological functionality, literature association, or computational recoverability.

### 4.4 Relationship to existing computational approaches

BioSensor-X builds on established computational strategies for regulatory inference and TF-activity estimation rather than introducing a fundamentally new statistical primitive. In *E. coli*, transcriptional regulation is organized through multitarget regulators, multifactor promoters, and hierarchical regulatory networks, providing important context for the structure of the regulatory evidence used in TF prioritization [15]. Network-inference methods such as ARACNE, TIGRESS, and tree-based approaches use different statistical assumptions to infer regulatory relationships from expression data [12,14,20]. Other frameworks, including SCENIC, ChEA3, decoupleR, and related approaches, integrate expression signatures with prior regulatory knowledge to estimate TF-associated activity or enrichment [1,13,16,22].

The methodological contribution of BioSensor-X lies instead in the explicit evaluation of heterogeneous evidence integration as a TF-ranking system [32]. The framework treats transcriptomic responsiveness, regulatory specificity, annotation density, and curated regulatory evidence as separable evidence dimensions, combines them using a prespecified score, and evaluates the resulting composite alongside individual evidence-based comparators. This design makes it possible to examine whether observed benchmark-associated performance arises from integration itself or is dominated by a particular underlying feature.

The present results illustrate why component-level evaluation is important. Although the composite score was designed to combine complementary information, its conditional AUROC of 0.5782 was lower than the 0.7945 achieved by the regulon-size/annotation-density comparator. The DEVELOPMENT analyses further showed that alternative integration formulations produced distinct ranking behavior rather than a uniformly superior solution. This does not imply that evidence integration is generally ineffective; rather, it demonstrates that the value of integration must be established empirically for the particular evidence sources, candidate universe, and benchmark being evaluated.

The distinction is also relevant to existing TF-activity methods. Differences in regulatory-resource coverage and prior network composition can influence inferred TF activity [10,22]. BioSensor-X complements these approaches by emphasizing the downstream evaluation of ranking behavior and its dependence on evidence structure, rather than assuming that adding additional evidence dimensions will necessarily improve prioritization.

### 4.5 Limitations and future directions

Several limitations define the scope of the present findings. First, the conditional LOCKED TEST evaluation included only five represented positive TFs, resulting in an uncertain AUROC estimate (95% bootstrap CI, 0.3308–0.8109). Second, candidate coverage was incomplete, with 5 of 12 LOCKED TEST TFs represented in the discovery candidate universe, and candidate generation drew contributed instances from only three of the six processed transcriptomic datasets. Third, the benchmark is not a direct experimental measure of biosensor functionality and may reflect differences in regulatory annotation and experimental attention. Fourth, Maximum aggregation was used as the prespecified TF-level aggregation strategy. The mean candidate-level score was retained descriptively but was not used for TF ranking. Finally, no independent experimental validation of newly prioritized TFs was performed, and the study was restricted to *E. coli* and the available regulatory and transcriptomic resources.

Future work should expand candidate coverage using additional biologically relevant *E. coli* datasets, evaluate the framework on independent benchmarks, and experimentally test prioritized TFs. More informative benchmarks should also distinguish experimentally demonstrated biosensor phenotypes from broader regulatory evidence and evaluate candidate coverage and ranking performance as separate endpoints. This would help determine whether transcriptomic responsiveness and other evidence dimensions provide reproducible information beyond annotation density when the endpoint is experimentally measured biosensor functionality.

Overall, the present findings support BioSensor-X as a leakage-controlled framework for evaluating heterogeneous TF-prioritization evidence, while highlighting the importance of candidate coverage and annotation structure when interpreting benchmark-associated performance.

## 5. Conclusion

BioSensor-X provides a leakage-controlled and coverage-aware framework for evaluating heterogeneous transcriptomic and regulatory evidence for transcription-factor (TF) prioritization. Under the corrected LOCKED TEST evaluation, the prespecified four-component score achieved a conditional AUROC of 0.5782 (95% TF-level bootstrap CI, 0.3308–0.8109) among the five benchmark-positive TFs represented in the discovery candidate universe. Candidate representation was incomplete, with 5 of 12 LOCKED TEST TFs covered by the discovery candidate space, corresponding to 41.7% candidate coverage.

A central finding was that the individual evidence dimensions behaved differently from the integrated score. The regulon-size/annotation-density comparator achieved an AUROC of 0.7945 (95% bootstrap CI, 0.6655–0.9055), exceeding the equal-weight composite. The accompanying annotation-density analyses showed that part of the observed benchmark-associated signal was related to the structure of curated regulatory information rather than providing direct evidence of biosensor functionality. Candidate coverage and conditional ranking were therefore treated as separate evaluation properties throughout the study.

The principal contribution of BioSensor-X is methodological: it provides a structured framework for evaluating evidence-integrated TF prioritization while explicitly separating candidate coverage, conditional ranking performance, and annotation-density-associated benchmark structure. The findings demonstrate that incorporating additional evidence dimensions does not necessarily improve benchmark discrimination and that component-level evaluation is essential before interpreting a composite score as evidence of biosensor suitability.

The present results should be interpreted within the scope of the *E. coli* discovery data and benchmark used here. Further evaluation on independent datasets, broader candidate spaces, and experimentally measured biosensor phenotypes will be required to determine whether the observed computational signals translate into reproducible evidence of biosensor functionality.

## 6. Statements and Declarations

### Data and Code Availability Statement

The datasets, regulatory resources, and benchmark information analyzed in this study were obtained from publicly accessible repositories and established *Escherichia coli* regulatory resources.

- **Transcriptomic Discovery Datasets** **NCBI Gene Expression Omnibus (GEO)** GSE4511: https://www.ncbi.nlm.nih.gov/geo/query/acc.cgi?acc=GSE4511 GSE4556: https://www.ncbi.nlm.nih.gov/geo/query/acc.cgi?acc=GSE4556 GSE12797: https://www.ncbi.nlm.nih.gov/geo/query/acc.cgi?acc=GSE12797 GSE20095: https://www.ncbi.nlm.nih.gov/geo/query/acc.cgi?acc=GSE20095 GSE34023: https://www.ncbi.nlm.nih.gov/geo/query/acc.cgi?acc=GSE34023 GSE80002: https://www.ncbi.nlm.nih.gov/geo/query/acc.cgi?acc=GSE80002
- **Regulatory Resources** **RegulonDB**: https://regulondb.ccg.unam.mx/ **EcoCyc**: https://ecocyc.org/ These resources were used to obtain *E. coli* transcription-factor target relationships, regulatory evidence, and regulon annotations.

### BioSensor-X Repository and Reproducibility Resources

The BioSensor-X repository contains the complete computational package used in this study, including the curated *E. coli* transcription-factor biosensor benchmark and its classifications, TF-family partitions, candidate-coverage metadata, processed candidate-level feature matrices, standardized evidence components, TF-level scores, benchmark evaluation outputs, comparator and sensitivity-analysis results, and all custom scripts used for transcriptomic preprocessing, TF-activity estimation, evidence-component construction, score calculation, candidate-to-TF aggregation, statistical analysis, and generation of the main and supplementary figures and tables. The repository also includes the associated parameter files, computational-environment information, and documentation required to reproduce the reported analyses.

### BioSensor-X Repository: https://github.com/mohdmehboobuddin/BioSensor-X

Large primary datasets obtained from external public repositories are not redistributed within the repository and should be downloaded directly from their original sources using the accession numbers and links provided above.

### CRediT Authorship Contribution Statement

### Mohd Mehboob Uddin

Conceptualization, Methodology, Software, Data Curation, Formal Analysis, Investigation, Visualization, Validation, Writing – Original Draft, Writing – Review & Editing.

### Syed Mohd Zakariya Ali Khan

Conceptualization, Methodology, Software, Data Curation, Formal Analysis, Investigation, Writing – Review & Editing.

### Sridhar Gunde

Conceptualization, Methodology, Formal Analysis, Investigation, Writing – Review & Editing.

### Declaration of Competing Interest

The authors declare that there are no known competing financial interests or personal relationships that could have appeared to influence the work reported in this study.

### Funding

This research did not receive any specific grant from funding agencies in the public, commercial, or not-for-profit sectors.

## Acknowledgements

The authors gratefully acknowledge the publicly available datasets and regulatory resources provided by the NCBI Gene Expression Omnibus (GEO), RegulonDB, and EcoCyc, which made this computational study possible.

## Notes

https://github.com/mohdmehboobuddin/BioSensor-X

### Competing Interest Statement

The authors have declared no competing interest.

