## Supplementary Information for "BioSensor-X: A Coverage-Aware Evidence-Integration and Benchmarking Framework for Transcription-Factor Prioritization in Microbial Biosensor Engineering"

**Supplementary Tables**

**Table S1. Per-TF Scores, Rankings, and Best-Candidate Characteristics**

Table S1 provides the complete TF-level output for the 60 transcription factors represented in the BioSensor-X candidate universe. The information is divided into four panels to facilitate presentation of the complete TF-level results.

**Table S1A. TF-Level Scores and Rankings**

Table S1A reports the BioSensor-X Primary score and TF-level ranking, together with the candidate context associated with the maximum Primary score, the number of candidate contexts, the mean candidate-level score, and corrected benchmark-membership indicators.

| **TF** | **BioSensor-X Primary Score** | **TF Rank** | **Best Candidate** | **n Candidates** | **Mean Candidate Score** | **DEVELOPMENT** | **LOCKED TEST** |
| --- | --- | --- | --- | --- | --- | --- | --- |
| PurR | 2.185510 | 1 | PurR_Aerobic_pH_8.7_vs_Aerobic_pH_7.0_GSE4511 | 6 | 0.596900 | 0 | 0 |
| TorR | 1.949503 | 2 | TorR_rpoS_vs_WT_Stat_GSE12797 | 6 | 0.418390 | 0 | 0 |
| Cra | 1.734516 | 3 | Cra_rpoS_vs_WT_Stat_GSE12797 | 6 | 0.601974 | 0 | 0 |
| FlhDC | 1.732949 | 4 | FlhDC_rpoS_vs_WT_Exp_GSE12797 | 6 | 0.071722 | 0 | 0 |
| PhoP | 1.578344 | 5 | PhoP_rpoS_vs_WT_Stat_GSE12797 | 6 | 0.403173 | 0 | 1 |
| Nac | 1.453492 | 6 | Nac_rpoS_vs_WT_Stat_GSE12797 | 6 | 0.639095 | 0 | 0 |
| ArgP | 1.419079 | 7 | ArgP_rpoS_vs_WT_Exp_GSE12797 | 6 | 0.494725 | 0 | 0 |
| IHF | 1.389013 | 8 | IHF_rpoS_vs_WT_Exp_GSE12797 | 6 | 0.551631 | 0 | 0 |
| SoxS | 1.351552 | 9 | SoxS_Aerobic_pH_5.0_vs_Aerobic_pH_7.0_GSE4511 | 6 | 0.635794 | 1 | 0 |
| CpxR | 1.336185 | 10 | CpxR_Aerobic_pH_5.0_vs_Aerobic_pH_7.0_GSE4511 | 6 | 0.677155 | 1 | 0 |
| Fis | 1.335155 | 11 | Fis_Aerobic_pH_5.0_vs_Aerobic_pH_7.0_GSE4511 | 6 | 0.286935 | 0 | 0 |
| ArcA | 1.313945 | 12 | ArcA_rpoS_vs_WT_Stat_GSE12797 | 6 | 0.249317 | 1 | 0 |
| PtrR | 1.261204 | 13 | PtrR_rpoS_vs_WT_Stat_GSE12797 | 6 | 0.128113 | 0 | 0 |
| Lrp | 1.222421 | 14 | Lrp_Aerobic_pH_5.0_vs_Aerobic_pH_7.0_GSE4511 | 6 | 0.950252 | 1 | 0 |
| MarA | 1.194585 | 15 | MarA_rpoS_vs_WT_Stat_GSE12797 | 6 | 0.635036 | 1 | 0 |
| CRP | 1.127690 | 16 | CRP_rpoS_vs_WT_Stat_GSE12797 | 6 | 0.442678 | 1 | 0 |
| MetR | 1.114470 | 17 | MetR_Aerobic_pH_8.7_vs_Aerobic_pH_7.0_GSE4511 | 6 | 0.035087 | 0 | 0 |
| GadE | 1.078530 | 18 | GadE_Anaerobic_pH_8.5_vs_Anaerobic_pH_7.0_GSE4556 | 6 | 0.575487 | 0 | 0 |
| CysB | 1.034118 | 19 | CysB_Anaerobic_pH_8.5_vs_Anaerobic_pH_7.0_GSE4556 | 6 | 0.285366 | 0 | 0 |
| ArgR | 1.019566 | 20 | ArgR_rpoS_vs_WT_Exp_GSE12797 | 6 | 0.497002 | 0 | 1 |
| OmpR | 0.992120 | 21 | OmpR_Aerobic_pH_5.0_vs_Aerobic_pH_7.0_GSE4511 | 6 | 0.164596 | 1 | 0 |
| LexA | 0.986377 | 22 | LexA_Aerobic_pH_8.7_vs_Aerobic_pH_7.0_GSE4511 | 6 | -0.142208 | 0 | 1 |
| LrhA | 0.977368 | 23 | LrhA_rpoS_vs_WT_Stat_GSE12797 | 6 | -0.071426 | 0 | 0 |
| NagC | 0.940238 | 24 | NagC_Anaerobic_pH_8.5_vs_Anaerobic_pH_7.0_GSE4556 | 6 | 0.322477 | 0 | 0 |
| NarL | 0.915566 | 25 | NarL_Anaerobic_pH_5.7_vs_Anaerobic_pH_7.0_GSE4556 | 6 | -0.005070 | 1 | 0 |
| RcdA | 0.901064 | 26 | RcdA_Aerobic_pH_5.0_vs_Aerobic_pH_7.0_GSE4511 | 6 | 0.321288 | 0 | 0 |
| MetJ | 0.899929 | 27 | MetJ_Anaerobic_pH_5.7_vs_Anaerobic_pH_7.0_GSE4556 | 6 | 0.129455 | 0 | 0 |
| H-NS | 0.884843 | 28 | H-NS_Anaerobic_pH_5.7_vs_Anaerobic_pH_7.0_GSE4556 | 6 | 0.232800 | 1 | 0 |
| SlyA | 0.874695 | 29 | SlyA_Anaerobic_pH_8.5_vs_Anaerobic_pH_7.0_GSE4556 | 6 | 0.194265 | 0 | 0 |
| OxyR | 0.868717 | 30 | OxyR_Anaerobic_pH_5.7_vs_Anaerobic_pH_7.0_GSE4556 | 6 | 0.562955 | 1 | 0 |
| Fur | 0.831719 | 31 | Fur_rpoS_vs_WT_Exp_GSE12797 | 6 | 0.328764 | 0 | 1 |
| TyrR | 0.801291 | 32 | TyrR_Aerobic_pH_5.0_vs_Aerobic_pH_7.0_GSE4511 | 6 | 0.080777 | 0 | 0 |
| YidZ | 0.708068 | 33 | YidZ_Aerobic_pH_8.7_vs_Aerobic_pH_7.0_GSE4511 | 6 | -0.383372 | 0 | 0 |
| YieP | 0.707493 | 34 | YieP_Anaerobic_pH_5.7_vs_Anaerobic_pH_7.0_GSE4556 | 6 | -0.389218 | 0 | 0 |
| FNR | 0.704271 | 35 | FNR_Anaerobic_pH_5.7_vs_Anaerobic_pH_7.0_GSE4556 | 6 | 0.230432 | 1 | 0 |
| GalR | 0.692488 | 36 | GalR_Aerobic_pH_5.0_vs_Aerobic_pH_7.0_GSE4511 | 6 | 0.083187 | 0 | 0 |
| PlaR | 0.687110 | 37 | PlaR_Anaerobic_pH_8.5_vs_Anaerobic_pH_7.0_GSE4556 | 6 | 0.075735 | 0 | 0 |
| BasR | 0.680161 | 38 | BasR_Anaerobic_pH_8.5_vs_Anaerobic_pH_7.0_GSE4556 | 6 | 0.097432 | 0 | 0 |
| GlaR | 0.677847 | 39 | GlaR_Aerobic_pH_5.0_vs_Aerobic_pH_7.0_GSE4511 | 6 | 0.357962 | 0 | 0 |
| FadR | 0.677319 | 40 | FadR_Anaerobic_pH_5.7_vs_Anaerobic_pH_7.0_GSE4556 | 6 | 0.322217 | 0 | 0 |
| GalS | 0.667440 | 41 | GalS_Aerobic_pH_5.0_vs_Aerobic_pH_7.0_GSE4511 | 6 | 0.066816 | 0 | 0 |
| PdhR | 0.653205 | 42 | PdhR_Aerobic_pH_8.7_vs_Aerobic_pH_7.0_GSE4511 | 6 | 0.320777 | 0 | 0 |
| Rob | 0.631984 | 43 | Rob_rpoS_vs_WT_Stat_GSE12797 | 6 | -0.107001 | 1 | 0 |
| TrpR | 0.576891 | 44 | TrpR_Aerobic_pH_5.0_vs_Aerobic_pH_7.0_GSE4511 | 6 | -0.042333 | 0 | 0 |
| DnaA | 0.563455 | 45 | DnaA_Aerobic_pH_8.7_vs_Aerobic_pH_7.0_GSE4511 | 6 | 0.154955 | 0 | 0 |
| NtrC | 0.507512 | 46 | NtrC_Aerobic_pH_8.7_vs_Aerobic_pH_7.0_GSE4511 | 6 | -0.253417 | 0 | 0 |
| GntR | 0.503696 | 47 | GntR_Aerobic_pH_8.7_vs_Aerobic_pH_7.0_GSE4511 | 6 | 0.020603 | 0 | 0 |
| Mlc | 0.494867 | 48 | Mlc_Aerobic_pH_8.7_vs_Aerobic_pH_7.0_GSE4511 | 6 | -0.322561 | 0 | 0 |
| AcrR | 0.440004 | 49 | AcrR_rpoS_vs_WT_Stat_GSE12797 | 6 | 0.092694 | 0 | 0 |
| ZraR | 0.427740 | 50 | ZraR_rpoS_vs_WT_Exp_GSE12797 | 6 | -0.123741 | 0 | 0 |
| AscG | 0.404247 | 51 | AscG_Aerobic_pH_8.7_vs_Aerobic_pH_7.0_GSE4511 | 6 | -0.060946 | 0 | 0 |
| CytR | 0.359710 | 52 | CytR_Anaerobic_pH_8.5_vs_Anaerobic_pH_7.0_GSE4556 | 6 | -0.222564 | 0 | 0 |
| PhoB | 0.304756 | 53 | PhoB_Anaerobic_pH_5.7_vs_Anaerobic_pH_7.0_GSE4556 | 6 | 0.077066 | 0 | 1 |
| YfeC | 0.275767 | 54 | YfeC_Anaerobic_pH_5.7_vs_Anaerobic_pH_7.0_GSE4556 | 6 | -0.363402 | 0 | 0 |
| YeiE | 0.271558 | 55 | YeiE_rpoS_vs_WT_Exp_GSE12797 | 6 | -0.107065 | 0 | 0 |
| RpoN | 0.264965 | 56 | RpoN_Aerobic_pH_5.0_vs_Aerobic_pH_7.0_GSE4511 | 6 | -0.459800 | 0 | 0 |
| NsrR | 0.220062 | 57 | NsrR_Anaerobic_pH_8.5_vs_Anaerobic_pH_7.0_GSE4556 | 6 | -0.047709 | 0 | 0 |
| LldR | 0.148575 | 58 | LldR_Aerobic_pH_8.7_vs_Aerobic_pH_7.0_GSE4511 | 6 | -0.449792 | 0 | 0 |
| DpiA | 0.097388 | 59 | DpiA_Anaerobic_pH_8.5_vs_Anaerobic_pH_7.0_GSE4556 | 6 | -0.432502 | 0 | 0 |
| SoxR | 0.077356 | 60 | SoxR_rpoS_vs_WT_Stat_GSE12797 | 6 | -0.294918 | 1 | 0 |

**Table S1B. Alternative Composite Scores and Benchmark Membership**

Table S1B reports the retained composite-score outputs and comparator values for each TF.

| **TF** | **Equal-weight Composite** | **Tier-weighted Composite** | **Ordinal Composite** | **B1 Expression Only** | **B2 Regulatory Only** | **B4 Custom TF-Activity Proxy** | **B7 Benchmark-Information Control** | **B8 Annotation-Density Diagnostic** |
| --- | --- | --- | --- | --- | --- | --- | --- | --- |
| PurR | 2.185510 | 2.310303 | 2.559888 | 3.494657 | 0.892816 | 6.204225 | 0 | 0.786830 |
| TorR | 1.949503 | 2.345339 | 3.137011 | 6.917913 | -0.315593 | 11.025881 | 0 | 0.367615 |
| Cra | 1.734516 | 1.782139 | 1.877386 | 1.951266 | 1.251143 | 4.030358 | 0 | 1.503484 |
| FlhDC | 1.732949 | 1.990485 | 2.505557 | 4.151921 | -0.081937 | 7.129980 | 0 | 0.834928 |
| PhoP | 1.578344 | 1.586390 | 1.602482 | 1.159718 | 1.022735 | 2.915463 | 1 | 1.046668 |
| Nac | 1.453492 | 1.527408 | 1.675242 | 0.880088 | 1.435238 | 1.957420 | 0 | 3.869278 |
| ArgP | 1.419079 | 1.526540 | 1.741463 | 3.148031 | 0.558538 | 5.716002 | 0 | 0.118275 |
| IHF | 1.389013 | 1.344926 | 1.256752 | 0.892987 | 1.500232 | 1.446879 | 0 | 2.001662 |
| SoxS | 1.351552 | 1.393604 | 1.477707 | 1.839831 | 1.094152 | 3.873402 | 1 | 1.189503 |
| CpxR | 1.336185 | 1.366180 | 1.426172 | 1.598715 | 1.126472 | 3.533790 | 1 | 1.254143 |
| Fis | 1.335155 | 1.465824 | 1.727161 | 1.614567 | 0.545143 | 3.556117 | 0 | 2.089087 |
| ArcA | 1.313945 | 1.493529 | 1.852696 | 2.592872 | 0.288008 | 4.934061 | 1 | 1.574816 |
| PtrR | 1.261204 | 1.566031 | 2.175685 | 5.097736 | -0.753609 | 8.462161 | 0 | -0.508417 |
| Lrp | 1.222421 | 1.347135 | 1.596563 | 1.495480 | 1.251210 | 3.388384 | 1 | 3.501221 |
| MarA | 1.194585 | 1.222539 | 1.278447 | 1.557884 | 1.041535 | 3.476279 | 1 | 1.084269 |
| CRP | 1.127690 | 1.193053 | 1.323777 | 0.835241 | 1.046039 | 1.716445 | 1 | 3.090880 |
| MetR | 1.114470 | 1.123898 | 1.142752 | 1.187346 | 0.245192 | 2.954376 | 0 | -0.508417 |
| GadE | 1.078530 | 1.150233 | 1.293640 | 2.432863 | 0.454800 | 4.708688 | 0 | -0.089201 |
| CysB | 1.034118 | 1.051870 | 1.087374 | 1.353843 | 0.509267 | 3.188888 | 0 | 0.019732 |
| ArgR | 1.019566 | 0.985472 | 0.917284 | 0.884478 | 1.263307 | 1.728387 | 1 | 1.527812 |
| OmpR | 0.992120 | 1.141508 | 1.440285 | 1.988963 | 0.172652 | 4.083454 | 1 | 1.344106 |
| LexA | 0.986377 | 1.068723 | 1.233416 | 0.868489 | 0.023933 | 2.194878 | 1 | 1.046668 |
| LrhA | 0.977368 | 0.945316 | 0.881214 | 0.806130 | 0.150940 | 1.785930 | 0 | -0.696921 |
| NagC | 0.940238 | 0.960853 | 1.002083 | 1.411097 | 0.509267 | 3.269530 | 0 | 0.019732 |
| NarL | 0.915566 | 1.031467 | 1.263268 | 1.319206 | 0.004423 | 3.140101 | 1 | 1.007647 |
| RcdA | 0.901064 | 0.935589 | 1.004638 | 1.689290 | 0.454800 | 3.661365 | 0 | -0.089201 |
| MetJ | 0.899929 | 0.901433 | 0.904440 | 1.028871 | 0.245192 | 2.731165 | 0 | -0.508417 |
| H-NS | 0.884843 | 0.979484 | 1.168767 | 0.894023 | 0.516416 | 2.541231 | 1 | 2.031633 |
| SlyA | 0.874695 | 0.883276 | 0.900438 | 1.170417 | 0.454800 | 2.930532 | 0 | -0.089201 |
| OxyR | 0.868717 | 0.844672 | 0.796583 | 0.678255 | 1.003224 | 2.011477 | 1 | 1.007647 |
| Fur | 0.831719 | 0.887242 | 0.998287 | 0.745092 | 0.899965 | 1.439260 | 1 | 2.798731 |
| TyrR | 0.801291 | 0.759888 | 0.677082 | 0.873217 | 0.509267 | 1.522490 | 0 | 0.019732 |
| YidZ | 0.708068 | 0.899124 | 1.281236 | 2.822319 | -0.847861 | 5.257237 | 0 | -0.696921 |
| YieP | 0.707493 | 0.826811 | 1.065447 | 1.387556 | -0.673920 | 3.236373 | 0 | -0.349038 |
| FNR | 0.704271 | 0.799552 | 0.990115 | 0.906826 | 0.428038 | 2.559264 | 1 | 1.854877 |
| GalR | 0.692488 | 0.664084 | 0.607277 | 0.714334 | 0.324882 | 1.888680 | 0 | -0.349038 |
| PlaR | 0.687110 | 0.638421 | 0.541042 | 0.696087 | 0.324882 | 1.317237 | 0 | -0.349038 |
| BasR | 0.680161 | 0.636694 | 0.549761 | 0.909871 | 0.454800 | 1.464362 | 0 | -0.089201 |
| GlaR | 0.677847 | 0.739637 | 0.863218 | 0.534782 | 0.652047 | 1.615827 | 0 | 2.302894 |
| FadR | 0.677319 | 0.637757 | 0.558634 | 0.803890 | 0.783578 | 1.574365 | 0 | 0.568355 |
| GalS | 0.667440 | 0.635042 | 0.570248 | 0.754235 | 0.324882 | 1.776178 | 0 | -0.349038 |
| PdhR | 0.653205 | 0.609310 | 0.521521 | 0.876286 | 0.841077 | 1.452297 | 0 | 0.683353 |
| Rob | 0.631984 | 0.759542 | 1.014657 | 1.552352 | -0.246563 | 3.468488 | 1 | 0.505675 |
| TrpR | 0.576891 | 0.520382 | 0.407364 | 0.897499 | 0.245192 | 1.096956 | 0 | -0.508417 |
| DnaA | 0.563455 | 0.518579 | 0.428826 | 0.827463 | 0.454800 | 1.424640 | 0 | -0.089201 |
| NtrC | 0.507512 | 0.568320 | 0.689936 | 0.775043 | -0.353904 | 1.588150 | 0 | 0.290994 |
| GntR | 0.503696 | 0.449290 | 0.340479 | 0.752226 | 0.245192 | 1.156202 | 0 | -0.508417 |
| Mlc | 0.494867 | 0.606912 | 0.831002 | 1.242097 | -0.673920 | 3.031494 | 0 | -0.349038 |
| AcrR | 0.440004 | 0.379298 | 0.257885 | 0.796865 | 0.393911 | 0.978721 | 0 | -0.210979 |
| ZraR | 0.427740 | 0.502237 | 0.651231 | 0.847084 | 0.004423 | 1.973767 | 0 | 1.007647 |
| AscG | 0.404247 | 0.354733 | 0.255707 | 0.848181 | 0.150940 | 1.294020 | 0 | -0.696921 |
| CytR | 0.359710 | 0.480666 | 0.722578 | 1.420315 | -0.489535 | 3.282513 | 0 | 0.019732 |
| PhoB | 0.304756 | 0.369357 | 0.498559 | 0.415909 | 0.042734 | 1.694994 | 1 | 1.084269 |
| YfeC | 0.275767 | 0.360119 | 0.528822 | 0.704860 | -0.673920 | 2.251373 | 0 | -0.349038 |
| YeiE | 0.271558 | 0.346220 | 0.495543 | 0.667760 | -0.246563 | 1.978409 | 0 | 0.505675 |
| RpoN | 0.264965 | 0.332677 | 0.468100 | 0.888220 | -0.673920 | 1.782625 | 0 | -0.349038 |
| NsrR | 0.220062 | 0.263135 | 0.349283 | 0.806888 | 0.078400 | 1.088574 | 0 | 1.155601 |
| LldR | 0.148575 | 0.196804 | 0.293261 | 0.879228 | -0.489535 | 1.233792 | 0 | 0.019732 |
| DpiA | 0.097388 | 0.158817 | 0.281676 | 0.734536 | -0.753609 | 1.605648 | 0 | -0.508417 |
| SoxR | 0.077356 | 0.146935 | 0.286091 | 0.392764 | -0.673920 | 1.835208 | 1 | -0.349038 |

**Table S1C. Best-Candidate Transcriptomic Characteristics and Discovery Context**

Table S1C reports the transcriptomic characteristics and discovery context associated with the highest-scoring candidate for each TF.

| **TF** | **GSE** | **Contrast** | **Signal Context** | **z_resp** | **mlm_z** | **mlm_p** | **P90_bg** | **n_bg** |
| --- | --- | --- | --- | --- | --- | --- | --- | --- |
| PurR | GSE4511 | Aerobic_pH_8.7_vs_Aerobic_pH_7.0 | high_pH_aerobic | 3.494630 | 6.204225 | 0.000000 | 1.510619 | 5 |
| TorR | GSE12797 | rpoS_vs_WT_Stat | rpoS_knockout_stationary | 6.917854 | -11.025881 | 0.000000 | 6.825419 | 5 |
| Cra | GSE12797 | rpoS_vs_WT_Stat | rpoS_knockout_stationary | 1.951252 | -4.030358 | 0.004000 | 1.519614 | 5 |
| FlhDC | GSE12797 | rpoS_vs_WT_Exp | rpoS_knockout_exponential | 4.151921 | 7.129980 | 0.000000 | 2.176769 | 5 |
| PhoP | GSE12797 | rpoS_vs_WT_Stat | rpoS_knockout_stationary | 1.159718 | -2.915463 | 0.018000 | 0.827398 | 5 |
| Nac | GSE12797 | rpoS_vs_WT_Stat | rpoS_knockout_stationary | 0.479537 | 1.957420 | 0.718000 | 0.745242 | 5 |
| ArgP | GSE12797 | rpoS_vs_WT_Exp | rpoS_knockout_exponential | 3.148031 | -5.716002 | 0.002000 | 3.742974 | 5 |
| IHF | GSE12797 | rpoS_vs_WT_Exp | rpoS_knockout_exponential | 0.117063 | 1.446879 | 0.117000 | 0.557552 | 5 |
| SoxS | GSE4511 | Aerobic_pH_5.0_vs_Aerobic_pH_7.0 | low_pH_aerobic | 1.839831 | 3.873402 | 0.000000 | 2.584740 | 5 |
| CpxR | GSE4511 | Aerobic_pH_5.0_vs_Aerobic_pH_7.0 | low_pH_aerobic | 1.598715 | -3.533790 | 0.006000 | 2.209830 | 5 |
| Fis | GSE4511 | Aerobic_pH_5.0_vs_Aerobic_pH_7.0 | low_pH_aerobic | 1.614567 | 3.556117 | 0.006000 | 1.249180 | 5 |
| ArcA | GSE12797 | rpoS_vs_WT_Stat | rpoS_knockout_stationary | 2.592872 | -4.934061 | 0.001000 | 2.256640 | 5 |
| PtrR | GSE12797 | rpoS_vs_WT_Stat | rpoS_knockout_stationary | 5.097736 | -8.462161 | 0.001000 | 5.407920 | 5 |
| Lrp | GSE4511 | Aerobic_pH_5.0_vs_Aerobic_pH_7.0 | low_pH_aerobic | 1.495480 | 3.388384 | 0.005000 | 3.039970 | 5 |
| MarA | GSE12797 | rpoS_vs_WT_Stat | rpoS_knockout_stationary | 1.557884 | -3.476279 | 0.009000 | 2.672980 | 5 |
| CRP | GSE12797 | rpoS_vs_WT_Stat | rpoS_knockout_stationary | 0.308447 | -1.716445 | 0.047000 | 0.775938 | 5 |
| MetR | GSE4511 | Aerobic_pH_8.7_vs_Aerobic_pH_7.0 | high_pH_aerobic | 1.187346 | 2.954376 | 0.019000 | 0.971156 | 5 |
| GadE | GSE4556 | Anaerobic_pH_8.5_vs_Anaerobic_pH_7.0 | high_pH_anaerobic | 2.432845 | 4.708688 | 0.000000 | 4.012983 | 5 |
| CysB | GSE4556 | Anaerobic_pH_8.5_vs_Anaerobic_pH_7.0 | high_pH_anaerobic | 1.353835 | 3.188888 | 0.004000 | 1.721489 | 5 |
| ArgR | GSE12797 | rpoS_vs_WT_Exp | rpoS_knockout_exponential | 0.316926 | -1.728387 | 0.123000 | 1.253679 | 5 |
| OmpR | GSE4511 | Aerobic_pH_5.0_vs_Aerobic_pH_7.0 | low_pH_aerobic | 1.988949 | -4.083454 | 0.002000 | 2.362999 | 5 |
| LexA | GSE4511 | Aerobic_pH_8.7_vs_Aerobic_pH_7.0 | high_pH_aerobic | 0.648119 | 2.194878 | 0.086000 | 0.585548 | 5 |
| LrhA | GSE12797 | rpoS_vs_WT_Stat | rpoS_knockout_stationary | 0.357779 | 1.785930 | 0.086000 | 0.476389 | 5 |
| NagC | GSE4556 | Anaerobic_pH_8.5_vs_Anaerobic_pH_7.0 | high_pH_anaerobic | 1.411089 | -3.269530 | 0.009000 | 2.241235 | 5 |
| NarL | GSE4556 | Anaerobic_pH_5.7_vs_Anaerobic_pH_7.0 | low_pH_anaerobic | 1.319198 | 3.140101 | 0.007000 | 1.271985 | 5 |
| RcdA | GSE4511 | Aerobic_pH_5.0_vs_Aerobic_pH_7.0 | low_pH_aerobic | 1.689279 | 3.661365 | 0.007000 | 3.054587 | 5 |
| MetJ | GSE4556 | Anaerobic_pH_5.7_vs_Anaerobic_pH_7.0 | low_pH_anaerobic | 1.028866 | 2.731165 | 0.014000 | 1.253148 | 5 |
| H-NS | GSE4556 | Anaerobic_pH_5.7_vs_Anaerobic_pH_7.0 | low_pH_anaerobic | 0.894019 | 2.541231 | 0.013000 | 1.487971 | 5 |
| SlyA | GSE4556 | Anaerobic_pH_8.5_vs_Anaerobic_pH_7.0 | high_pH_anaerobic | 1.170411 | 2.930532 | 0.008000 | 1.910768 | 5 |
| OxyR | GSE4556 | Anaerobic_pH_5.7_vs_Anaerobic_pH_7.0 | low_pH_anaerobic | 0.517910 | 2.011477 | 0.052000 | 1.737554 | 5 |
| Fur | GSE12797 | rpoS_vs_WT_Exp | rpoS_knockout_exponential | 0.111654 | 1.439260 | 0.077000 | 0.940275 | 5 |
| TyrR | GSE4511 | Aerobic_pH_5.0_vs_Aerobic_pH_7.0 | low_pH_aerobic | 0.170745 | 1.522490 | 0.126000 | 0.721636 | 5 |
| YidZ | GSE4511 | Aerobic_pH_8.7_vs_Aerobic_pH_7.0 | high_pH_aerobic | 2.822298 | -5.257237 | 0.000000 | 2.927524 | 5 |
| YieP | GSE4556 | Anaerobic_pH_5.7_vs_Anaerobic_pH_7.0 | low_pH_anaerobic | 1.387548 | 3.236373 | 0.004000 | 1.058966 | 5 |
| FNR | GSE4556 | Anaerobic_pH_5.7_vs_Anaerobic_pH_7.0 | low_pH_anaerobic | 0.906822 | -2.559264 | 0.012000 | 2.070972 | 5 |
| GalR | GSE4511 | Aerobic_pH_5.0_vs_Aerobic_pH_7.0 | low_pH_aerobic | 0.430728 | 1.888680 | 0.059000 | 1.060885 | 5 |
| PlaR | GSE4556 | Anaerobic_pH_8.5_vs_Anaerobic_pH_7.0 | high_pH_anaerobic | 0.025022 | -1.317237 | 0.209000 | 0.606453 | 5 |
| BasR | GSE4556 | Anaerobic_pH_8.5_vs_Anaerobic_pH_7.0 | high_pH_anaerobic | 0.129475 | 1.464362 | 0.114000 | 0.826030 | 5 |
| GlaR | GSE4511 | Aerobic_pH_5.0_vs_Aerobic_pH_7.0 | low_pH_aerobic | 0.237011 | 1.615827 | 0.142000 | 1.217986 | 5 |
| FadR | GSE4556 | Anaerobic_pH_5.7_vs_Anaerobic_pH_7.0 | low_pH_anaerobic | 0.207574 | -1.574365 | 0.117000 | 1.373953 | 5 |
| GalS | GSE4511 | Aerobic_pH_5.0_vs_Aerobic_pH_7.0 | low_pH_aerobic | 0.350855 | 1.776178 | 0.084000 | 1.008635 | 5 |
| PdhR | GSE4511 | Aerobic_pH_8.7_vs_Aerobic_pH_7.0 | high_pH_aerobic | 0.120910 | 1.452297 | 0.206000 | 1.375373 | 5 |
| Rob | GSE12797 | rpoS_vs_WT_Stat | rpoS_knockout_stationary | 1.552343 | -3.468488 | 0.008000 | 2.198743 | 5 |
| TrpR | GSE4511 | Aerobic_pH_5.0_vs_Aerobic_pH_7.0 | low_pH_aerobic | -0.131371 | 1.096956 | 0.201000 | 0.538051 | 5 |
| DnaA | GSE4511 | Aerobic_pH_8.7_vs_Aerobic_pH_7.0 | high_pH_aerobic | 0.101275 | 1.424640 | 0.132000 | 1.028343 | 5 |
| NtrC | GSE4511 | Aerobic_pH_8.7_vs_Aerobic_pH_7.0 | high_pH_aerobic | 0.217362 | 1.588150 | 0.118000 | 0.588729 | 5 |
| GntR | GSE4511 | Aerobic_pH_8.7_vs_Aerobic_pH_7.0 | high_pH_aerobic | -0.089308 | -1.156202 | 0.145000 | 0.676553 | 5 |
| Mlc | GSE4511 | Aerobic_pH_8.7_vs_Aerobic_pH_7.0 | high_pH_aerobic | 1.242090 | 3.031494 | 0.017000 | 1.387646 | 5 |
| AcrR | GSE12797 | rpoS_vs_WT_Stat | rpoS_knockout_stationary | -0.215314 | 0.978721 | 0.161000 | 0.730154 | 5 |
| ZraR | GSE12797 | rpoS_vs_WT_Exp | rpoS_knockout_exponential | 0.491138 | 1.973767 | 0.048000 | 1.451993 | 5 |
| AscG | GSE4511 | Aerobic_pH_8.7_vs_Aerobic_pH_7.0 | high_pH_aerobic | 0.008538 | -1.294020 | 0.122000 | 0.899864 | 5 |
| CytR | GSE4556 | Anaerobic_pH_8.5_vs_Anaerobic_pH_7.0 | high_pH_anaerobic | 1.420306 | -3.282513 | 0.004000 | 2.751960 | 5 |
| PhoB | GSE4556 | Anaerobic_pH_5.7_vs_Anaerobic_pH_7.0 | low_pH_anaerobic | 0.293217 | 1.694994 | 0.096000 | 1.572910 | 5 |
| YfeC | GSE4556 | Anaerobic_pH_5.7_vs_Anaerobic_pH_7.0 | low_pH_anaerobic | 0.688229 | -2.251373 | 0.027000 | 1.216293 | 5 |
| YeiE | GSE12797 | rpoS_vs_WT_Exp | rpoS_knockout_exponential | 0.494433 | -1.978409 | 0.071000 | 1.570804 | 5 |
| RpoN | GSE4511 | Aerobic_pH_5.0_vs_Aerobic_pH_7.0 | low_pH_aerobic | 0.355433 | -1.782625 | 0.061000 | 0.829577 | 5 |
| NsrR | GSE4556 | Anaerobic_pH_8.5_vs_Anaerobic_pH_7.0 | high_pH_anaerobic | -0.137322 | -1.088574 | 0.348000 | 0.996650 | 5 |
| LldR | GSE4511 | Aerobic_pH_8.7_vs_Aerobic_pH_7.0 | high_pH_aerobic | -0.034222 | 1.233792 | 0.203000 | 0.724362 | 5 |
| DpiA | GSE4556 | Anaerobic_pH_8.5_vs_Anaerobic_pH_7.0 | high_pH_anaerobic | 0.229784 | -1.605648 | 0.115000 | 0.912853 | 5 |
| SoxR | GSE12797 | rpoS_vs_WT_Stat | rpoS_knockout_stationary | 0.392765 | -1.835208 | 0.042000 | 1.307987 | 5 |

**Table S1D. Best-Candidate Regulatory and Annotation Characteristics**

Table S1D reports the regulatory and annotation characteristics associated with the same highest-scoring candidate.

| **TF** | **z_spec** | **z_mech_canonical** | **z_reg** | **n_targets** | **Highest Tier** | **Perturbation Class** |
| --- | --- | --- | --- | --- | --- | --- |
| PurR | 3.461751 | 0.786830 | 0.998801 | 20 | Tier A (STRONG) | pH |
| TorR | 1.511287 | 0.367615 | -0.998801 | 13 | Tier B (MODERATE) | TF_perturbation |
| Cra | 2.484514 | 1.503484 | 0.998801 | 41 | Tier A (STRONG) | TF_perturbation |
| FlhDC | 2.943747 | 0.834928 | -0.998801 | 21 | Tier B (MODERATE) | TF_perturbation |
| PhoP | 3.108191 | 1.046668 | 0.998801 | 26 | Tier A (STRONG) | TF_perturbation |
| Nac | 2.463958 | 3.869278 | -0.998801 | 413 | Tier B (MODERATE) | TF_perturbation |
| ArgP | 1.411209 | 0.118275 | 0.998801 | 10 | Tier A (STRONG) | TF_perturbation |
| IHF | 2.438529 | 2.001662 | 0.998801 | 67 | Tier A (STRONG) | TF_perturbation |
| SoxS | 1.378074 | 1.189503 | 0.998801 | 30 | Tier A (STRONG) | pH |
| CpxR | 1.493079 | 1.254143 | 0.998801 | 32 | Tier A (STRONG) | pH |
| Fis | 2.635770 | 2.089087 | -0.998801 | 73 | Tier B (MODERATE) | pH |
| ArcA | 2.086893 | 1.574816 | -0.998801 | 44 | Tier B (MODERATE) | TF_perturbation |
| PtrR | 1.454299 | -0.508417 | -0.998801 | 5 | Tier B (MODERATE) | TF_perturbation |
| Lrp | 0.891783 | 3.501221 | -0.998801 | 289 | Tier B (MODERATE) | pH |
| MarA | 1.137386 | 1.084269 | 0.998801 | 27 | Tier A (STRONG) | TF_perturbation |
| CRP | 2.110237 | 3.090880 | -0.998801 | 194 | Tier B (MODERATE) | TF_perturbation |
| MetR | 2.780151 | -0.508417 | 0.998801 | 5 | Tier A (STRONG) | pH |
| GadE | 0.971658 | -0.089201 | 0.998801 | 8 | Tier A (STRONG) | pH |
| CysB | 1.764095 | 0.019732 | 0.998801 | 9 | Tier A (STRONG) | pH |
| ArgR | 1.234725 | 1.527812 | 0.998801 | 42 | Tier A (STRONG) | TF_perturbation |
| OmpR | 1.634213 | 1.344106 | -0.998801 | 35 | Tier B (MODERATE) | pH |
| LexA | 3.249523 | 1.046668 | -0.998801 | 26 | Tier B (MODERATE) | pH |
| LrhA | 3.249812 | -0.696921 | 0.998801 | 4 | Tier A (STRONG) | TF_perturbation |
| NagC | 1.331322 | 0.019732 | 0.998801 | 9 | Tier A (STRONG) | pH |
| NarL | 2.334214 | 1.007647 | -0.998801 | 25 | Tier B (MODERATE) | pH |
| RcdA | 1.005367 | -0.089201 | 0.998801 | 8 | Tier A (STRONG) | pH |
| MetJ | 2.080462 | -0.508417 | 0.998801 | 5 | Tier A (STRONG) | pH |
| H-NS | 1.612519 | 2.031633 | -0.998801 | 69 | Tier B (MODERATE) | pH |
| SlyA | 1.418765 | -0.089201 | 0.998801 | 8 | Tier A (STRONG) | pH |
| OxyR | 0.950507 | 1.007647 | 0.998801 | 25 | Tier A (STRONG) | pH |
| Fur | 1.415296 | 2.798731 | -0.998801 | 146 | Tier B (MODERATE) | TF_perturbation |
| TyrR | 2.015888 | 0.019732 | 0.998801 | 9 | Tier A (STRONG) | pH |
| YidZ | 1.705675 | -0.696921 | -0.998801 | 4 | Tier B (MODERATE) | pH |
| YieP | 2.790257 | -0.349038 | -0.998801 | 6 | Tier B (MODERATE) | pH |
| FNR | 1.054182 | 1.854877 | -0.998801 | 58 | Tier B (MODERATE) | pH |
| GalR | 1.689462 | -0.349038 | 0.998801 | 6 | Tier A (STRONG) | pH |
| PlaR | 2.073660 | -0.349038 | 0.998801 | 6 | Tier A (STRONG) | pH |
| BasR | 1.681571 | -0.089201 | 0.998801 | 8 | Tier A (STRONG) | pH |
| GlaR | 1.170285 | 2.302894 | -0.998801 | 90 | Tier B (MODERATE) | pH |
| FadR | 0.934546 | 0.568355 | 0.998801 | 16 | Tier A (STRONG) | pH |
| GalS | 1.669142 | -0.349038 | 0.998801 | 6 | Tier A (STRONG) | pH |
| PdhR | 0.809758 | 0.683353 | 0.998801 | 18 | Tier A (STRONG) | pH |
| Rob | 1.468711 | 0.505675 | -0.998801 | 15 | Tier B (MODERATE) | TF_perturbation |
| TrpR | 1.948554 | -0.508417 | 0.998801 | 5 | Tier A (STRONG) | pH |
| DnaA | 1.242950 | -0.089201 | 0.998801 | 8 | Tier A (STRONG) | pH |
| NtrC | 2.520495 | 0.290994 | -0.998801 | 12 | Tier B (MODERATE) | pH |
| GntR | 1.613713 | -0.508417 | 0.998801 | 5 | Tier A (STRONG) | pH |
| Mlc | 2.085212 | -0.349038 | -0.998801 | 6 | Tier B (MODERATE) | pH |
| AcrR | 1.187512 | -0.210979 | 0.998801 | 7 | Tier A (STRONG) | TF_perturbation |
| ZraR | 1.210977 | 1.007647 | -0.998801 | 25 | Tier B (MODERATE) | TF_perturbation |
| AscG | 1.306573 | -0.696921 | 0.998801 | 4 | Tier A (STRONG) | pH |
| CytR | 0.997596 | 0.019732 | -0.998801 | 9 | Tier B (MODERATE) | pH |
| PhoB | 0.840342 | 1.084269 | -0.998801 | 27 | Tier B (MODERATE) | pH |
| YfeC | 1.762676 | -0.349038 | -0.998801 | 6 | Tier B (MODERATE) | pH |
| YeiE | 1.084926 | 0.505675 | -0.998801 | 15 | Tier B (MODERATE) | TF_perturbation |
| RpoN | 2.052268 | -0.349038 | -0.998801 | 6 | Tier B (MODERATE) | pH |
| NsrR | 0.860775 | 1.155601 | -0.998801 | 29 | Tier B (MODERATE) | pH |
| LldR | 1.607596 | 0.019732 | -0.998801 | 9 | Tier B (MODERATE) | pH |
| DpiA | 1.666989 | -0.508417 | -0.998801 | 5 | Tier B (MODERATE) | pH |
| SoxR | 1.264501 | -0.349038 | -0.998801 | 6 | Tier B (MODERATE) | TF_perturbation |

Each TF was represented by six candidate contexts, and the TF-level BioSensor-X Primary score was obtained by maximum aggregation across the corresponding candidate contexts.

**Table S1A note:** Each TF was represented by six candidate contexts. The TF-level BioSensor-X Primary score was obtained by maximum aggregation across those candidate contexts, with rank 1 corresponding to the highest score. Benchmark-membership indicators use the corrected mutually exclusive TF-level DEVELOPMENT and LOCKED TEST partitions.

**Table S1B note:** BioSensor-X Primary and benchmark-membership status are reported in Table S1A. B1 Expression Only, B2 Regulatory Only, and B4 Custom TF-Activity Proxy are comparator scores. B7 is the benchmark-information control defined by membership in the DEVELOPMENT or LOCKED TEST benchmark union and is not a biological evidence component. B8 is the annotation-density diagnostic based on canonical Tier A+B regulon target count (n_targets). Composite and comparator values are reported from the final canonical analysis outputs.

**Table S1C note:** Each row reports the transcriptomic characteristics and discovery context of the highest-scoring candidate identified in Table S1A. z_resp denotes the standardized transcriptomic responsiveness component. mlm_z is the TF-associated enrichment statistic derived from the permutation-based analysis of limma moderated t-statistics, and mlm_p is its empirical P value.

**Table S1D note:** Each row reports the regulatory and annotation characteristics of the highest-scoring candidate identified in Table S1A. z_spec denotes standardized regulatory specificity, z_mech_canonical denotes the standardized canonical regulon-size/annotation-density component, and z_reg denotes the standardized curated regulatory-evidence component. n_targets represents the canonical Tier A+B regulatory target count used in the annotation-density component.

**Table S2. Extended Performance Comparisons and Evaluation Population**

Table S2 summarizes the performance of BioSensor-X and the retained comparator methods on the LOCKED TEST evaluation set, together with ranking metrics and the coverage of the benchmark within the candidate universe. Because the candidate universe contains only five of the twelve TFs in the full LOCKED TEST benchmark, discrimination and ranking metrics are conditional on the five represented positive TFs and the 55 negative TFs in the candidate universe. Candidate coverage is reported separately to distinguish candidate-generation representation from conditional ranking performance.

**Table S2A. Discrimination Performance**

**Table S2A.** Discrimination performance of BioSensor-X and retained comparator methods on the conditional LOCKED TEST evaluation set. AUROC values were evaluated using the five represented LOCKED TEST TFs and 55 negative TFs. Bootstrap confidence intervals were calculated at the TF level. Top-5, Top-10, and Top-20 values denote the number of represented positive TFs recovered within the corresponding ranks, with five represented positive TFs available for evaluation. B7 is a benchmark-information control based on benchmark-union membership rather than a biological scoring component. B8 is an annotation-density diagnostic based on curated regulatory target count.

| **Method** | **AUROC** | **95% Bootstrap CI** | **Top-5** | **Top-10** | **Top-20** | **Role** |
| --- | --- | --- | --- | --- | --- | --- |
| **BioSensor-X Primary** | **0.5782** | **0.3308–0.8109** | 1/5 | 1/5 | 2/5 | Primary (equal-weight) |
| B1 Expression Only | 0.2945 | 0.1127–0.4909 | 0/5 | 0/5 | 0/5 | Comparator |
| B2 Regulatory Only | 0.6545 | 0.3818–0.9091 | 1/5 | 2/5 | 3/5 | Comparator |
| B4 Custom TF-Activity Proxy | 0.3527 | 0.1964–0.5236 | 0/5 | 0/5 | 0/5 | Comparator |
| B7 Benchmark-Information Control | 0.8818 | 0.8273–0.9364 | 0/5 | 5/5 | 5/5 | Benchmark-information control |
| B8 Annotation-Density Diagnostic | 0.7945 | 0.6655–0.9055 | 1/5 | 1/5 | 5/5 | Annotation-density diagnostic |

**Table S2B. Ranking and Statistical Comparison**

**Table S2B.** Ranking performance of BioSensor-X and retained comparator methods on the conditional LOCKED TEST evaluation set. MRR denotes mean reciprocal rank calculated from the ranks of the five represented positive TFs, and median rank denotes the median rank of these represented positive TFs among the 60 TFs in the candidate universe. The bootstrap confidence interval column reports the corresponding TF-level AUROC confidence interval shown in Table S2A and is provided as the discrimination-performance uncertainty associated with each method.

| **Method** | **MRR** | **Median Rank** | **95% Bootstrap CI (AUROC)** | **Role** |
| --- | --- | --- | --- | --- |
| **BioSensor-X Primary** | **0.069316** | **22.0** | **0.3308–0.8109** | Primary (equal-weight) |
| B1 Expression Only | 0.026014 | 39.0 | 0.1127–0.4909 | Comparator |
| B2 Regulatory Only | 0.112746 | 12.0 | 0.3818–0.9091 | Comparator |
| B4 Custom TF-Activity Proxy | 0.027519 | 41.0 | 0.1964–0.5236 | Comparator |
| B7 Benchmark-Information Control | 0.105263 | 9.5 | 0.8273–0.9364 | Benchmark-information control |
| B8 Annotation-Density Diagnostic | 0.099819 | 18.0 | 0.6655–0.9055 | Annotation-density diagnostic |

**Table S2C. Evaluation Population and Candidate Coverage**

**Table S2C.** Composition of the LOCKED TEST benchmark and its representation within the BioSensor-X candidate universe. The full LOCKED TEST benchmark contains 12 TFs, of which five are represented in the 60-TF candidate universe. The remaining seven benchmark TFs are absent from the candidate universe and therefore cannot contribute to conditional ranking metrics. Candidate coverage is calculated as the proportion of the full benchmark represented in the candidate universe.

| **Full LOCKED TEST Benchmark TFs** | **Represented in Candidate Universe** | **Unrepresented in Candidate Universe** | **Candidate Coverage** | **Represented TFs** | **Unrepresented TFs** | **Conditional Evaluation Positive TFs** | **Conditional Evaluation Negative TFs** |
| --- | --- | --- | --- | --- | --- | --- | --- |
| 12 | 5 | 7 | 41.7% | ArgR, Fur, LexA, PhoB, PhoP | AraC, BgaR, CueR, EnvZ, EvgA, MerR, PhoQ | 5 | 55 |

The LOCKED TEST benchmark comprises 12 predefined transcription factors, whereas the BioSensor-X candidate universe contains 60 TFs. Five LOCKED TEST TFs, ArgR, Fur, LexA, PhoB, and PhoP, are represented in the candidate universe, giving a candidate coverage of 5/12 (41.7%). The seven remaining benchmark TFs, AraC, BgaR, CueR, EnvZ, EvgA, MerR, and PhoQ, are not represented and therefore cannot be ranked by BioSensor-X or by the retained comparator methods within the candidate universe.

Accordingly, conditional benchmark performance was evaluated using the five represented positive TFs and the remaining 55 TFs in the candidate universe as negatives. This distinction separates **candidate coverage**, which reflects representation of the external benchmark within the candidate-generation stage, from **conditional ranking performance**, which evaluates discrimination and rank ordering among TFs that are present in the candidate universe.

The primary BioSensor-X score achieved an AUROC of 0.5782 (95% bootstrap CI, 0.3308–0.8109), with one of five represented positives recovered within the Top-5 and Top-10 and two of five within the Top-20. The regulatory-only comparator B2 achieved an AUROC of 0.6545 (95% bootstrap CI, 0.3818–0.9091), while the expression-only comparator B1 and the custom TF-activity proxy B4 produced lower AUROC values of 0.2945 and 0.3527, respectively. The annotation-density diagnostic B8 achieved an AUROC of 0.7945 (95% bootstrap CI, 0.6655–0.9055), indicating that curated regulon size is strongly associated with benchmark membership within this candidate universe and should therefore be interpreted as an annotation-structure diagnostic rather than as direct evidence of biosensor suitability.

The benchmark-information control B7 achieved an AUROC of 0.8818 (95% bootstrap CI, 0.8273–0.9364). Because B7 is defined from benchmark-union membership, it is included as a control for the influence of benchmark-related information rather than as an independent biological predictor. Its performance therefore provides a reference for the degree to which benchmark-associated information can separate represented positives from the remaining candidate-universe TFs.

Together, these analyses show that conditional benchmark performance should be interpreted jointly with candidate coverage. The presence of only five benchmark positives within the candidate universe limits the scope of the conditional evaluation and emphasizes the importance of evaluating candidate-generation procedures for both ranking quality and representation of the broader biological search space.

**Table S3. Sensitivity and Stratified Analyses on the DEVELOPMENT Partition**

Table S3 summarizes predefined sensitivity and diagnostic analyses conducted on the DEVELOPMENT partition. These analyses were used to examine the behavior of the individual BioSensor-X evidence components, alternative component combinations, hierarchical formulations, residualized components, and regulon-size-stratified performance. All analyses were conducted independently of the LOCKED TEST endpoint and were not used to modify, tune, or select the final BioSensor-X Primary score.

The four panels provide complementary diagnostic views of the evidence framework. Table S3A reports component-subset sensitivity analyses, Table S3B reports hierarchical formulations, Table S3C reports residualized component analyses, and Table S3D reports analyses stratified by canonical regulon size. The DEVELOPMENT partition comprised 13 benchmark-positive TFs and 47 non-benchmark TFs within the 60-TF candidate universe.

**Table S3A. Component-Subset Sensitivity**

**Table S3A.** Component-subset sensitivity analyses on the DEVELOPMENT partition. AUROC values quantify discrimination of DEVELOPMENT benchmark membership using the indicated component subset. The delta is calculated relative to the z_mech-only reference. The four-component Primary corresponds to the prespecified equal-weight combination of z_resp, z_spec, z_mech, and z_reg.

| **Subset** | **Components** | **n_components** | **AUROC_DEVELOPMENT** | **Delta_vs_z_mech** |
| --- | --- | --- | --- | --- |
| z_mech | z_mech | 1 | 0.8044 | 0.0000 |
| z_resp | z_resp | 1 | 0.6923 | -0.1121 |
| z_spec | z_spec | 1 | 0.3863 | -0.4182 |
| z_reg | z_reg | 1 | 0.3773 | -0.4272 |
| z_mech+z_resp | z_mech,z_resp | 2 | 0.7709 | -0.0336 |
| z_mech+z_spec | z_mech,z_spec | 2 | 0.7185 | -0.0859 |
| z_mech+z_reg | z_mech,z_reg | 2 | 0.6416 | -0.1628 |
| z_resp+z_spec | z_resp,z_spec | 2 | 0.5532 | -0.2512 |
| z_resp+z_reg | z_resp,z_reg | 2 | 0.4730 | -0.3314 |
| z_spec+z_reg | z_spec,z_reg | 2 | 0.3142 | -0.4902 |
| z_mech+z_resp+z_spec | z_mech,z_resp,z_spec | 3 | 0.7005 | -0.1039 |
| z_mech+z_resp+z_reg | z_mech,z_resp,z_reg | 3 | 0.6809 | -0.1236 |
| z_mech+z_spec+z_reg | z_mech,z_spec,z_reg | 3 | 0.6039 | -0.2005 |
| z_resp+z_spec+z_reg | z_resp,z_spec,z_reg | 3 | 0.4223 | -0.3822 |
| **Primary (equal-weight 4-comp)** | **z_mech,z_resp,z_spec,z_reg** | **4** | **0.6285** | **-0.1759** |

**Table S3A note.** AUROC values were calculated on the DEVELOPMENT partition comprising 13 benchmark-positive TFs and 47 non-benchmark TFs. The z_mech component was used as the reference for the reported AUROC differences. Negative values indicate lower AUROC than the z_mech-only formulation. The Primary is the prespecified equal-weight combination of the four evidence components and was not selected on the basis of the DEVELOPMENT results.

**Table S3B. Hierarchical Models**

**Table S3B.** Hierarchical and gated formulations evaluated on the DEVELOPMENT partition. Delta values are calculated relative to the equal-weight four-component BioSensor-X Primary formulation. Gated models were evaluated using the predefined z_mech-based conditions indicated in the table.

| **Model** | **Gate** | **n_components** | **AUROC_DEVELOPMENT** | **Delta_vs_Primary** |
| --- | --- | --- | --- | --- |
| Primary (equal-weight 4-comp) | none | 4 | 0.6285 | 0.0000 |
| z_mech only | none | 1 | 0.8044 | +0.1759 |
| z_mech+z_resp (z_mech>0 gate) | z_mech > 0 | 2 | 0.7067 | +0.0782 |
| z_mech+z_reg (z_mech>0 gate) | z_mech > 0 | 2 | 0.6000 | -0.0285 |
| z_mech+z_resp+z_spec (z_mech>0 gate) | z_mech > 0 | 3 | 0.6200 | -0.0085 |
| z_mech+z_resp+z_reg (z_mech>0 gate) | z_mech > 0 | 3 | 0.6067 | -0.0218 |
| z_resp (low tertile gate) | z_mech in low tertile | 1 | 0.5000 | -0.1285 |
| z_mech (high tertile gate) | z_mech in high tertile | 1 | 0.5404 | -0.0881 |
| z_resp (high tertile gate) | z_mech in high tertile | 1 | 0.8081 | +0.1796 |

**Table S3B note.** The hierarchical formulations were evaluated on the DEVELOPMENT partition using the predefined component definitions and z_mech-based conditions shown above. These analyses were diagnostic and did not alter the primary LOCKED TEST analysis. Tertile-gated results characterize component behavior within the corresponding regulon-size strata rather than defining alternative versions of the final BioSensor-X score.

**Table S3C. Residualized Components**

**Table S3C.** Residualized component analyses on the DEVELOPMENT partition. Each non-annotation component was residualized with respect to z_mech to characterize benchmark-associated signal remaining after accounting for regulon-size/annotation-density structure. Combined residualized formulations were evaluated using the corresponding residualized components.

| **Residualized Component** | **AUROC_DEVELOPMENT** | **Delta_vs_z_mech** |
| --- | --- | --- |
| z_resp_resid | 0.6972 | -0.1072 |
| z_spec_resid | 0.3879 | -0.4165 |
| z_reg_resid | 0.5123 | -0.2921 |
| Primary_resid | 0.4910 | -0.3134 |
| z_resp_resid+z_spec_resid | 0.5646 | -0.2398 |
| z_resp_resid+z_reg_resid | 0.5385 | -0.2660 |
| z_resp_resid+z_spec_resid+z_reg_resid | 0.4910 | -0.3134 |

**Table S3C note.** Residualization was performed with respect to the regulon-size/annotation-density component z_mech. The resulting AUROCs characterize benchmark-associated signal remaining in the other evidence dimensions after accounting for annotation-density structure. Residualized analyses are diagnostic and should not be interpreted as evidence of causal or mechanistically independent effects.

**Table S3D. Regulon-Size-Stratified Analyses**

**Table S3D.** DEVELOPMENT-partition analyses stratified by canonical regulon size, defined by the number of curated Tier A+B regulatory targets. Tertile boundaries were determined from the 60-TF candidate universe using the canonical n_targets values and were not defined from LOCKED TEST performance.

| **Tertile** | **n_targets_range** | **n_TFs** | **n_positive_DEVELOPMENT** | **z_mech_AUROC** | **z_resp_AUROC** | **z_spec_AUROC** | **z_reg_AUROC** |
| --- | --- | --- | --- | --- | --- | --- | --- |
| Low | 4–8 | 23 | 1 | 0.5682 | 0.5000 | 0.1818 | 0.1591 |
| Medium | 9–25 | 17 | 3 | 0.8452 | 0.5714 | 0.4762 | 0.4167 |
| High | 26–413 | 20 | 9 | 0.5404 | 0.8081 | 0.3535 | 0.4848 |
| All | 4–413 | 60 | 13 | 0.8044 | 0.6923 | 0.3863 | 0.3773 |

**Table S3D note.** Analyses were stratified according to canonical n_targets, representing curated Tier A+B regulatory target count. The low, medium, and high strata contain 23, 17, and 20 TFs, respectively. The number of DEVELOPMENT positives varies across strata, and the low stratum contains only one positive TF; stratified AUROC values are therefore interpreted descriptively. Tertile boundaries were defined across the complete 60-TF candidate universe rather than from benchmark performance.

**Overall Table S3 Note**

All sensitivity, hierarchical, residualized, and stratified analyses were conducted on the DEVELOPMENT partition using the predefined BioSensor-X component definitions. The DEVELOPMENT partition comprised **13 benchmark-positive TFs and 47 non-benchmark TFs** within the 60-TF candidate universe. These analyses were diagnostic and were not selected or modified on the basis of LOCKED TEST results. The final LOCKED TEST BioSensor-X Primary score and endpoint were therefore unaffected by the sensitivity analyses.

**Supplementary Figures**

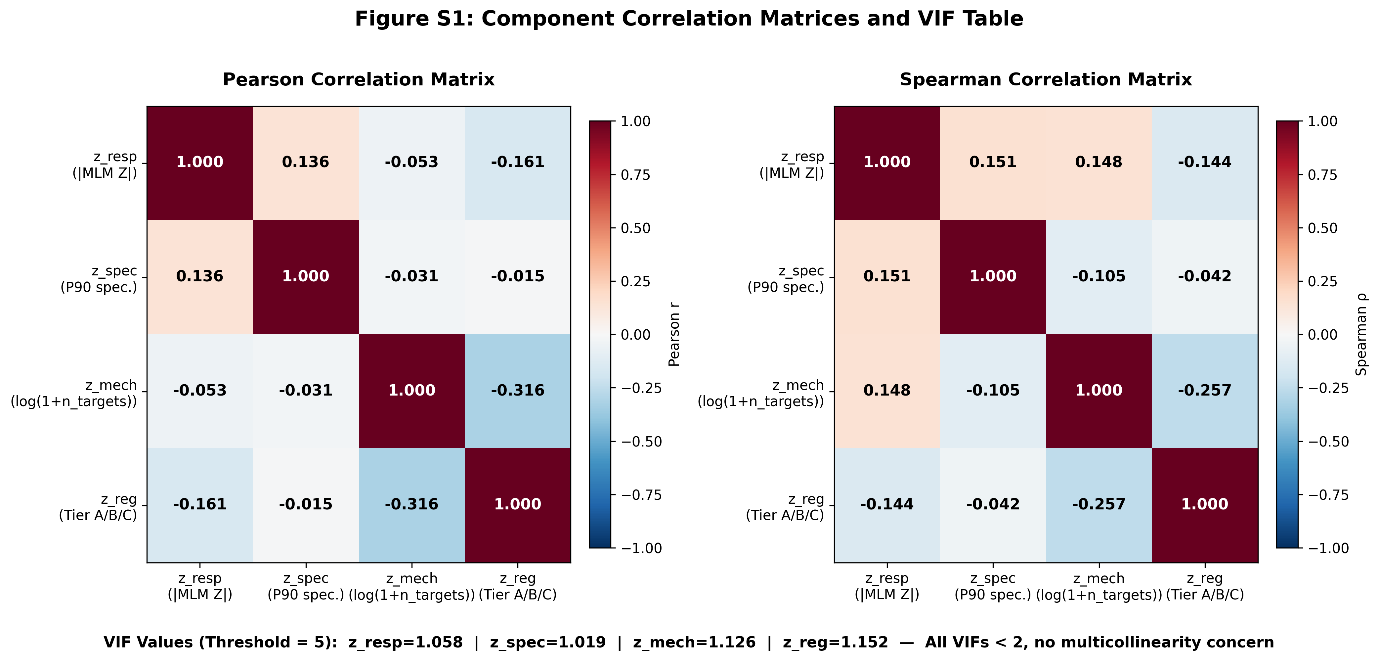

**Figure S1. Correlation Structure and Variance Inflation Diagnostics of BioSensor-X Evidence Components**

Figure S1 summarizes the relationships among the four evidence components used in the BioSensor-X Primary score.

The figure presents the component correlation structure together with variance-inflation diagnostics and pairwise relationships among the evaluated evidence dimensions.

**Figure S1 note:** The VIF diagnostics indicate low multicollinearity among the evaluated evidence components.

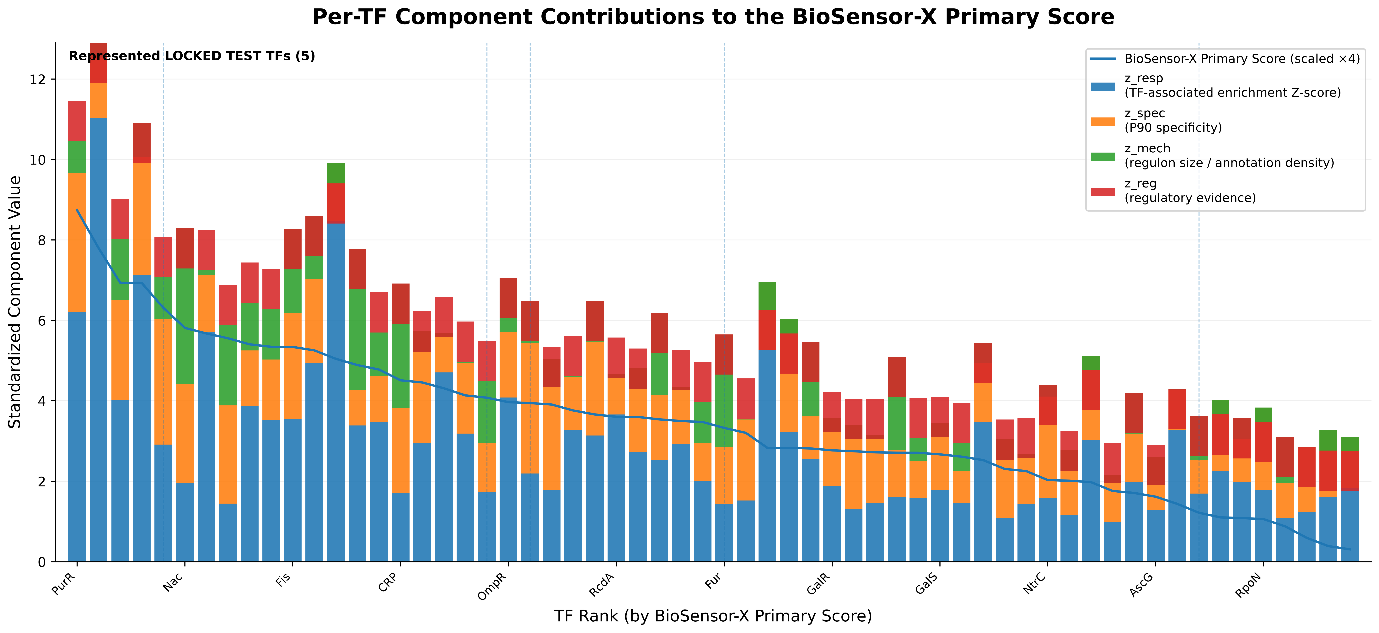

**Figure S2. Per-TF Component Contributions to the BioSensor-X Primary Score**

Figure S2 displays the contribution of the four standardized evidence components to the BioSensor-X Primary score for the 60 TFs in the candidate universe.

TFs are ordered according to their BioSensor-X TF-level ranking. The figure shows the relative contribution of transcriptional responsiveness, regulatory specificity, regulon size/annotation density, and curated regulatory evidence to the equal-weight composite score.

**Figure S2 note:** Component contributions are shown using the standardized evidence values used in construction of the BioSensor-X Primary score.

**Supplementary Note S1. Interpretation of the Regulon-Size/Annotation-Density Component**

The regulon-size component is derived from the number of curated regulatory targets associated with each TF and is therefore interpreted as an annotation-density/regulon-size proxy. This component should not be interpreted as a direct measure of mechanistic importance, regulatory quality, or biosensor suitability. Its benchmark-associated signal is considered in the context of the depth and distribution of prior regulatory annotation across the candidate TF population.

**Supplementary Note S2. Interpretation of Conditional Benchmark Metrics**

The LOCKED TEST analysis is conditional on representation within the discovery candidate universe. The corrected LOCKED TEST benchmark contains 12 TFs, of which 5 are represented in the candidate universe, corresponding to 41.7% candidate coverage. The five represented TFs are ArgR, Fur, LexA, PhoB, and PhoP. The seven unrepresented TFs are AraC, BgaR, CueR, EnvZ, EvgA, MerR, and PhoQ. TFs absent from the candidate universe cannot receive candidate-level scores and therefore do not contribute to the conditional discrimination analysis. Candidate coverage and conditional ranking performance should consequently be interpreted as distinct aspects of the evaluation.

**Supplementary Note S3. Relationship to the Main Manuscript**

The Supplementary Information provides expanded numerical outputs and diagnostic analyses supporting the principal findings reported in the main manuscript. The definitions of the BioSensor-X evidence components, scoring framework, benchmark construction, statistical procedures, and information-flow controls are described in the main Materials and Methods section and are not repeated here. The supplementary results should therefore be interpreted together with the corresponding descriptions and limitations presented in the main manuscript.
